# Drug targeting to sites of oxidative stress using the Baeyer-Villiger reaction

**DOI:** 10.1101/2021.09.03.458872

**Authors:** Thomas D. Avery, Jiahe Li, Dion J. L. Turner, Fisher R. Cherry, Mohd S. Ur Rasheed, Clarissa Aguilar, Andrew J. Shepherd, Jingxian Yu, Peter M. Grace, Andrew D. Abell

**Affiliations:** ARC Centre of Excellence for Nanoscale BioPhotonics (CNBP), Institute for Photonics and Advanced Sensing, Department of Chemistry, The University of Adelaide, Australia; Laboratories of Neuroimmunology, Department of Symptom Research, and the MD Anderson Pain Research Consortium, University of Texas MD Anderson Cancer Center, Houston, USA

**Author notes:** **Correspondence:** Andrew D. Abell, PhD, E, Peter M. Grace, PhD, E. Authors contributed equally (Thomas D. Avery, Jiahe Li).

**Keywords:** nerve injury, chemotherapy-induced peripheral neuropathy

## Abstract

The antioxidant nuclear factor erythroid 2-related factor 2 (Nrf2) is a desirable therapeutic target for a broad range of pathologies, including chronic diseases of the lung and liver, and autoimmune, neurodegenerative, and cardiovascular disorders. However, current Nrf2 activators are limited by unwanted effects due to non-specificity, and systemic distribution and action. Here we report that a 1,2-dicarbonyl moiety masks the electrophilic reactivity of the Nrf2 activator monomethyl fumarate (MMF), otherwise responsible for its non-specific effects. The 1,2-dicarbonyl compound is highly susceptible to Baeyer-Villiger oxidation, with generation of MMF specifically on exposure to pathological levels of hydrogen peroxide or peroxynitrite. Oral treatment with the MMF generating 1,2-dicarbonyl compound reversed chronic neuropathic and osteoarthritis pain in mice, and selectively activated Nrf2 at sites of oxidative stress. This 1,2-dicarbonyl platform may be used to treat additional disorders of oxidative stress, and to selectively target other therapeutics to sites of redox imbalance.

## INTRODUCTION

The ubiquity of oxidative stress as a pathological process has driven recent therapeutic interest in the antioxidant “master regulator” nuclear factor erythroid 2-related factor 2 (Nrf2)^1-3^. In preclinical models, small molecule Nrf2 activators alleviate neurodegenerative, cardiovascular, and other diseases, but unwanted effects have hampered clinical translation^2-4^. For example, prolonged Nrf2 activation with long-term systemic treatments can cause kidney and heart damage^5,6^. While electrophilic compounds activate Nrf2 by succinating thiols of Kelch like ECH-associated protein 1 (Keap1)—a protein that otherwise sequesters Nrf2 in the cytosol—such agents are also prone to adverse effects due to indiscriminate interactions with other protein thiols^1-4^.

We aimed to overcome these long-standing issues by creating a compound that would mask the unwanted electrophilic nature of the Nrf2 activator monomethyl fumarate (MMF) until selective generation of MMF at sites of oxidative pathology. We reasoned that these features would be conferred to MMF with the addition of a 1,2-dicarbonyl functional group; 1,2-dicarbonyl compounds are biologically compatible and substrates for Baeyer-Villiger oxidation by hydrogen peroxide or peroxynitrite^7-9^. Peroxides are an ideal trigger for localized MMF generation as their diffusion beyond sites of pathology is restricted by their inherent reactivity^3,10^. Here, we report realization of a MMF generating 1,2-dicarbonyl compound. After chemical synthesis and characterization, we used mouse models of chronic pain—conditions driven in part by localized overproduction of hydrogen peroxide and peroxynitrite^10,11^—to demonstrate therapeutic efficacy and preferential activation of Nrf2 at local sites of oxidative stress by this first-in-class oxidative pathology-activated drug.

## RESULTS

### Synthesis of MMF releasing 1,2-dicarbonyl compounds

As MMF is a carboxylic acid, we explored 1,2-diketones, α-keto amides and α-keto esters as 1,2-dicarbonyl masking groups. These biologically compatible functional groups are highly susceptible to the desired Baeyer-Villiger oxidation^7,9^, selectively reacting with hydrogen peroxide and peroxynitrite to form an anhydride that will hydrolyse, generating only MMF and acid **3** (Fig. 1a). Representatives of each 1,2-dicarbonyl chemotype (1,2-diketone (**1a**), α-keto amide (**1b**), α-keto ester (**1c**)) were synthesised (Fig. 1a; Fig. S1a-c).

**Figure 1.**
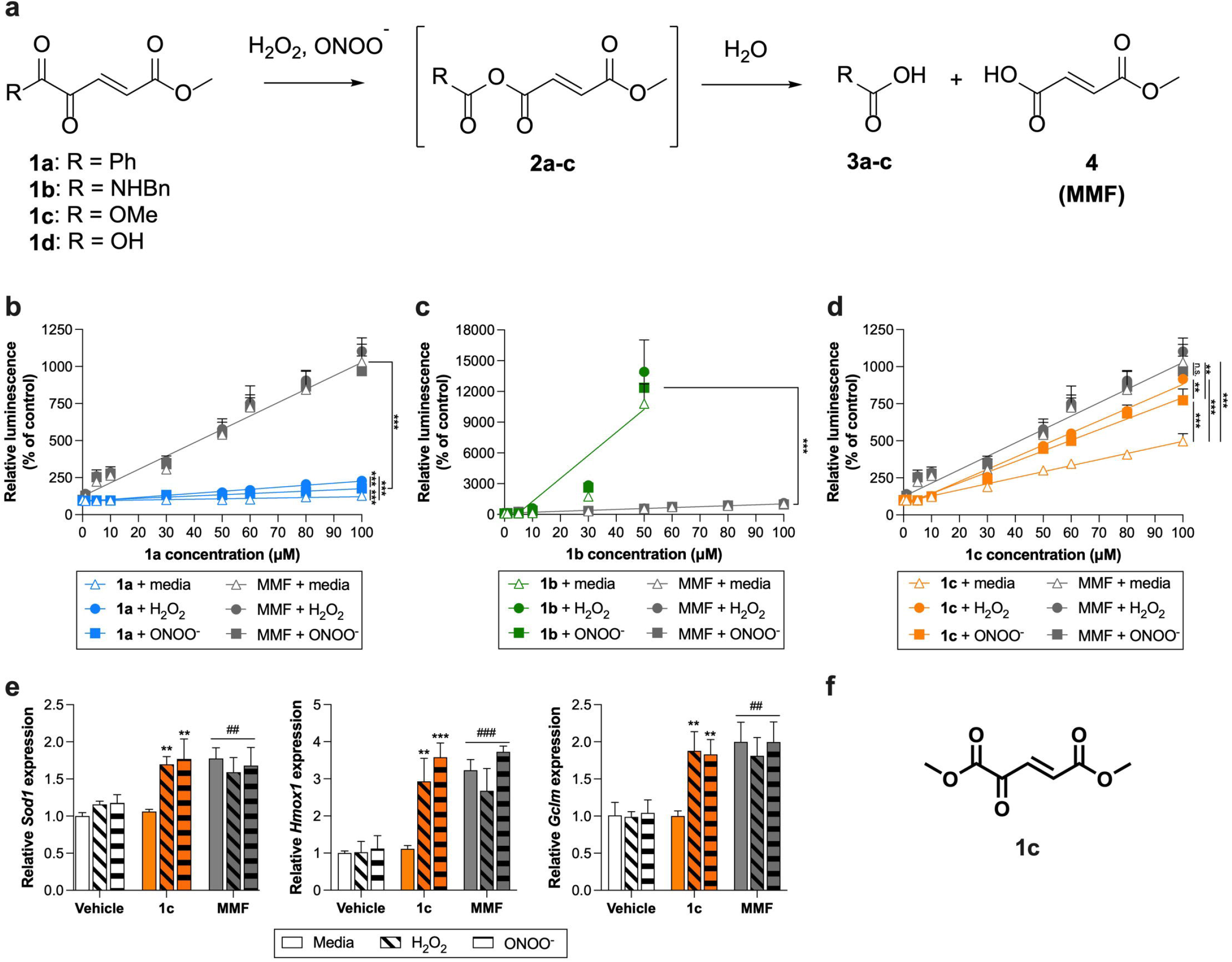
*In vitro* characterization of 1,2-dicarbonyl compounds. **(a)** MMF generating 1,2-dicarbonyl compounds and their reaction with aqueous hydrogen peroxide and peroxynitrite. **(b-e)** NRF2/antioxidant response element luciferase reporter HEK293 cells were treated with media, H_2_O_2_ (10 μM), or ONOO^-^ (20 μM). ARE-luciferase activity was quantified following treatment with concentration ranges of MMF or 1,2-dicarbonyl compounds **(b) 1a, (c) 1b**, or **(d) 1c**; not significant (n.s.), \*\**P* < 0.01, \*\*\**P* < 0.001. **(e)** Expression of Nrf2 target genes (*Sod1, Hmox1*, and *Gclm*) was quantified after treatment with **1c** or MMF (20 μM); relative to media: \*\**P* < 0.01, \*\*\**P* < 0.001; relative to vehicle: ^##^*P* < 0.01, ^###^*P* < 0.001. **(f)** Chemical structure of 1,2-dicarbonyl **1c**.

### MMF generation from 1,2-dicarbonyl compounds by peroxides

Each chemotype representative was evaluated by HPLC for its ability to generate MMF on exposure to peroxides. Compounds **1a-c** completely and quantitatively generated MMF within 30 min at 1 mM concentration in 100 mM phosphate buffer (pH7.4) containing five molar excess hydrogen peroxide (Fig. S2a-c). Both **1b** and **1c** were rapidly hydrolysed to α-keto acid **1d** (Fig. S2b,c), a 1,2-dicarbonyl compound also capable of undergoing the same Baeyer-Villiger like oxidative cleavage to generate MMF^12^. **1d** was synthesized, isolated, characterized, and underwent rapid conversion to MMF in hydrogen peroxide containing buffer (Fig. S2d). Compounds **1b-d** exhibit broad peaks in the HPLC due to equilibration with their hydrated form (geminal diol). By contrast, **1a** exists predominantly in its keto form under these conditions as determined by NMR, and as such has a much narrower peak shape in the HPLC traces (Supplementary Information; Tables S1-5).

The ability of the 1,2-dicarbonyl functional group to mask the electrophilicity and selectively generate MMF on exposure to peroxides in biological systems will be governed by a range of factors, including reaction rates with peroxides in cytosolic and extracellular fluids, and hydrolysis rates and the extent of hydration (geminal diol). Additional pharmacokinetic factors (e.g., membrane permeability and metabolism of the equilibrium components (Tables S1-6)) will also influence selective Nrf2 activation in the presence of peroxides. To gain initial insights into behaviour of the compounds in biological systems, we evaluated 1,2-dicarbonyl compounds **1a-c** for enhanced Nrf2 activity in the presence of pathological peroxide concentrations. Nrf2 activity was measured using a reporter HEK293 cell assay in which luciferase transcription is under the control of antioxidant response elements, themselves activated by Nrf2^1,2^. The ability of these compounds to activate Nrf2 was evaluated *in vitro*, with comparison made to MMF alone. As expected^13,14^, MMF increased Nrf2 activity (*P* < 0.001) and target gene expression above media control (*P* < 0.001), without enhancement by pathological concentrations of hydrogen peroxide or peroxynitrite^15,16^ (Fig. 1b-e).

Nrf2 activity induced by compound **1a** in physiological media was substantially lower than MMF (Fig. 1b; *P* < 0.001), consistent with reduced electrophilic reactivity. Although Nrf2 activity was enhanced by pathological concentrations of hydrogen peroxide or peroxynitrite (*P* < 0.001), activity remained substantially lower than that induced by MMF (Fig. 1b; *P* < 0.001). This suggests that **1a** converts poorly to MMF under the experimental conditions. In contrast, Nrf2 activity induced by compound **1b** was significantly higher than MMF, indicating greater electrophilic reactivity. However, hydrogen peroxide or peroxynitrite did not selectively enhance Nrf2 activation (*P* = 0.288), and **1b** was cytotoxic at higher concentrations (Fig. 1c; data not shown). This indicates that **1b** is more electrophilic than MMF and converts poorly to MMF under experimental conditions. We predict that with these unwanted features, compound **1b** would be prone to non-selective interactions with protein thiols, causing adverse effects.

Compound **1c** selectively enhanced Nrf2 activity in the presence of pathological concentrations of hydrogen peroxide (10 μM) and peroxynitrite (20 μM) (Fig. 1d; *P* < 0.001), but not physiological concentrations of hydrogen peroxide (1 μM) (Fig. S3a; *P* = 0.088) or peroxynitrite (2 μM) (Fig. S3b; *P* = 0.061)^15,16^. This suggests that **1c** is less electrophilic than MMF and has the appropriate rate of MMF production under the experimental conditions to selectively enhance Nrf2 activity. Increased Nrf2 activity was accompanied by elevated expression of target genes *Sod1* (*P* = 0.004), *Hmox1* (*P* = 0.001), and *Gclm* (*P* = 0.002) (Fig. 1e). We next tested whether the biological activity of **1c** was driven by MMF using the 1,2-dicarbonyl α-keto ester compound **5b** (dimethyl 2-oxoglutarate) (Fig. S1a). Compound **5b** is chemically similar to **1c** in its reactivity to hydrogen peroxide (Table S5) and susceptibility to hydrolysis (Table S6), also generating metabolites methanol and carbon dioxide in addition to non-electrophilic, monomethyl succinate (saturated MMF). Monomethyl succinate lacks the αβ-unsaturated system of MMF and thus cannot succinate Keap1 thiols, a key step in activation of Nrf2 by MMF. Compound **5b** failed to activate Nrf2 (Fig. S3c); with MMF being the only product absent after exposure of **5b** to peroxides, this result critically confirms MMF as the therapeutic metabolite of **1c**. Given selective activation of Nrf2 (on par with MMF) in the presence of pathological hydrogen peroxide and peroxynitrite, compound **1c** (Fig. 1f) was selected for validation in mouse models of chronic pain.

### 1,2-Dicarbonyl 1c reverses chronic pain

Peripheral nerve injury can lead to neuropathic pain^17^, driven in part by localised increases in hydrogen peroxide and peroxynitrite that hyperexcite sensory neurons^10,11^. We therefore examined the anti-nociceptive efficacy of 1,2-dicarbonyl compound **1c** in the spared nerve injury (SNI) model of neuropathic pain^18^ (Fig. 2a). Multi-day oral administration of compound **1c**, beginning at peak SNI-induced pain (postoperative day 7), dose-dependently reversed mechanical allodynia (Fig. 2b, Fig. S4a, *P* < 0.001) and dynamic allodynia as measured by changes in paw withdrawal responses (Fig. 2c, Fig. S4b, *P* < 0.001). Diroximel fumarate, FDA-approved prodrug that systemically distributes MMF^4^, also attenuated mechanical (Fig. 2b, *P* < 0.001) and dynamic allodynia (Fig. 2c, *P* < 0.001) at equi-mole doses. Treatment with compound **5b**, which generates saturated (non-electrophilic) MMF on exposure to peroxides, did not reverse SNI-induced allodynia (Fig. S4c, *P* = 0.864). This result confirms MMF as the therapeutic metabolite of **1c** *in vivo*. In sham-operated animals, the highest effective dose of compound **1c** did not alter withdrawal responses to mechanical (Fig. S4a, *P* = 0.139) or dynamic stimuli (Fig. S4b, *P* = 0.188) compared to vehicle, suggesting that **1c** does not alter normal sensory function. Dosing **1c** for five consecutive days did not lead to tolerance or a loss of efficacy, since reversal of mechanical allodynia increased over time (Fig. 2d, *P* < 0.001). In contrast, anti-nociceptive effects of morphine diminished when given over a similar timecourse (Fig. 2d, *P* = 0.025).

**Figure 2.**
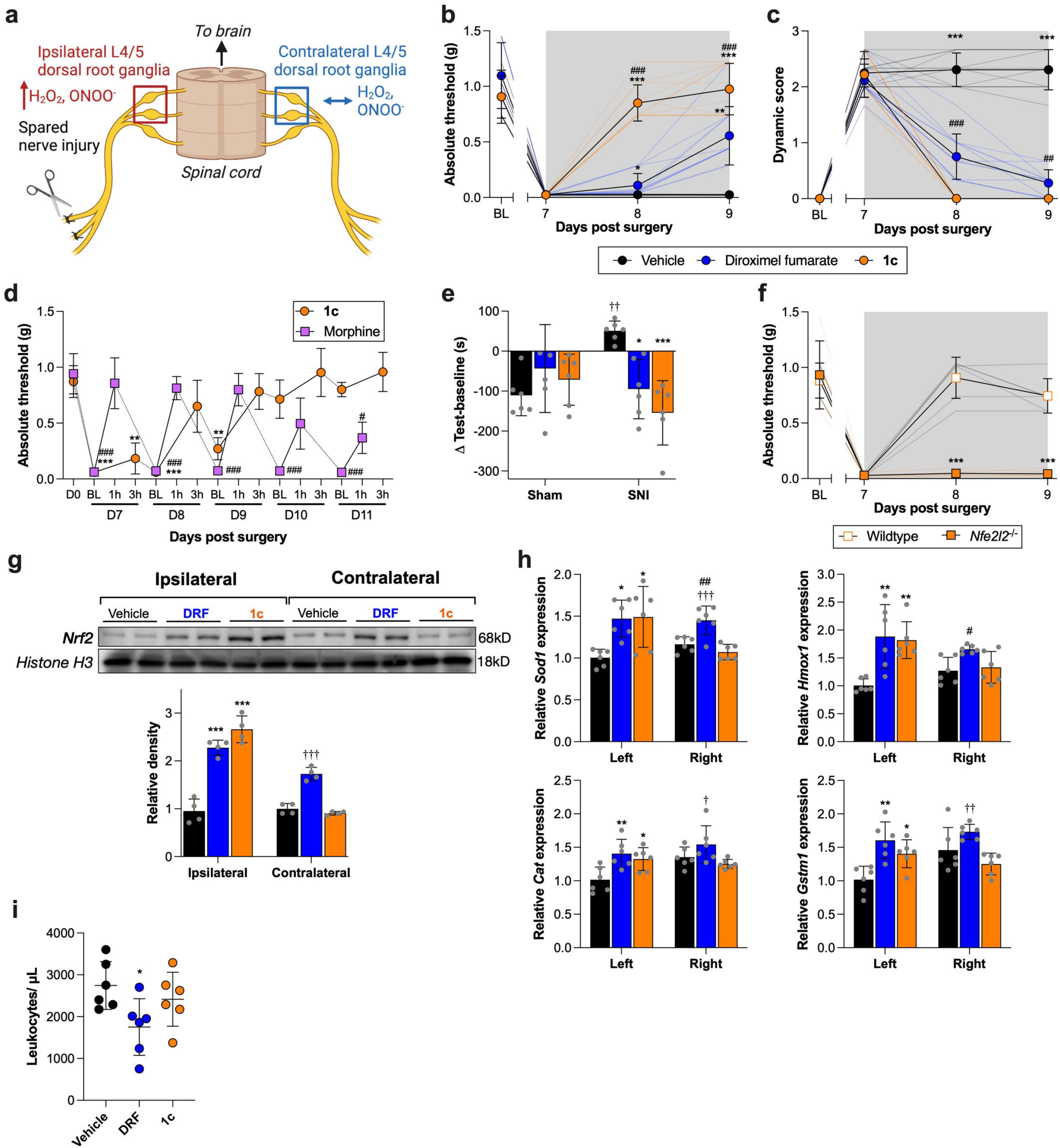
*In vivo* assessment of 1c efficacy and site-selectivity. **(a)** Schematic of spared nerve injury and tissues of interest. **(b, c)** Once neuropathic pain was established on day 7 after SNI, male (n=6) and female (n=6) mice were treated with oral **1c** (350 μmol/kg/day), diroximel fumarate (350 μmol/kg/day), or vehicle every day for 3 days (gray box). **(b)** Mechanical allodynia and **(c)** dynamic allodynia were assessed. Relative to vehicle: \**P* <0.05, \*\**P* <0.01, \*\*\**P* <0.001; **1c** vs. diroximel fumarate: ^##^*P* <0.01, ^###^*P* <0.001. **(d)** Seven days after SNI, male (n=3) and female (n=3) mice were treated with oral **1c** (350 μmol/kg/day) or subcutaneous morphine (3 mg/kg, b.i.d.). Relative to **1c** Day 0 (D0, prior to surgery): \*\**P* <0.01, \*\*\**P* <0.001; relative to morphine D0: ^#^*P* <0.05, ^###^*P* <0.001. **(e)** Male (n=3) and female (n=3) mice were treated with oral **1c** (350 µmol/kg/day), diroximel fumarate (350 µmol/kg/day) or vehicle every day for 7 days, beginning 7 days after SNI or sham surgery. Presence of ongoing pain was interpreted as an increase in time spent in retigabine-paired chamber (conditioning stimulus) after 7 days of treatment, relative to baseline. Relative to sham-vehicle: ^††^*P* <0.01; relative to SNI-vehicle: \**P* <0.05, \*\*\**P* <0.001. **(f)** Male and female *Nfe2l2*^-/-^ and wildtype control mice (n=3/sex/group) were treated with oral **1c** (350 μmol/kg/day), beginning 7 days after SNI, and mechanical allodynia assessed. Relative to wildtype controls: \*\*\**P* <0.001. **(g)** L4/5 DRG from 3 mice were pooled after 3 days of treatment, and nuclear extracts were probed for Nrf2 (n=2 males, n=2 females). Relative to ipsilateral vehicle: \*\*\**P* <0.001; relative to contralateral vehicle and **1c**: ^†††^*P* <0.001. **(h)** Antioxidant gene expression L4/5 DRG was determined after 3 days of treatment (n=3 males, n=3 females). Relative to ipsilateral vehicle: \**P* <0.05, \*\**P* <0.01; relative to contralateral vehicle: ^#^*P* <0.05; relative to contralateral **1c**: ^†^*P* <0.05; ^††^*P* <0.01, ^†††^*P* <0.001. (**i**) Naive male (n=3) and female (n = 3) mice were orally treated with **1c** (350 µmol/kg), diroximel fumarate (350 µmol/kg), or vehicle daily for 10 days. Blood was collected by cardiac puncture and leukocytes were manually counted. Relative to vehicle: \**P* <0.05. Individual replicates are presented in spaghetti plots or grey dots.

We further tested antinociceptive efficacy of compound **1c** in a clinically relevant models of cisplatin chemotherapy-induced peripheral neuropathy (CIPN)^19,20^ and osteoarthritis (surgical destabilization of the medial meniscus (DMM)^21^). Oxidative/nitrative species mediate in part cisplatin-CIPN and osteoarthritis pain^22,23^. When administered after the second cycle of cisplatin was complete, multi-day treatment with oral **1c** reversed mechanical allodynia (Fig. S4d, *P* < 0.001). We also assessed the effect of **1c** on numbness, a common symptom of CIPN^20^, by placing an adhesive patch on the hind paw and recording the latency to attend to the patch^19^. Mice treated with **1c** after the second cycle of cisplatin was complete had a faster response time (Fig. S4e, *P* = 0.011), indicating reversed numbness compared to mice treated with vehicle. Treatment with compound **1c** further reversed allodynia in the DMM model (Fig. S4f, *P* = 0.003). These data demonstrate anti-nociceptive efficacy of compound **1c** in preclinical chronic pain models of differing aetiologies, but with common underlying mechanisms of oxidative/nitrative stress.

In addition to stimulus-evoked pain, patients with chronic pain frequently experience ongoing (spontaneous/paroxysmal) pain^17^. We therefore assessed ongoing neuropathic pain using the conditioned place preference paradigm^24^, with the nerve blocker retigabine as the conditioning stimulus^19,25^. SNI-operated mice treated with vehicle spent more time in the retigabine-paired chamber, compared to sham-operated mice, indicating ongoing pain (Fig. 2e, *P* = 0.007). Compare the vehicle treated SNI-operated mice, those treated with compound **1c** had reduced time spent in the retigabine-paired chamber, indicating that compound **1c** had reversed ongoing pain (Fig. 2e, *P* < 0.001). Equi-mole diroximel fumarate treatment similarly reversed ongoing pain (Fig. 2e, *P* = 0.018).

### Site selective Nrf2 activation by 1,2-dicarbonyl 1c

Nrf2 activation was monitored as a proxy for site specific generation of MMF from compound **1c** in the SNI model. We first validated Nrf2 as a therapeutically-relevant biomarker by confirming that the anti-nociceptive effects of compound **1c** were dependent on Nrf2, as for other MMF-releasing compounds^26^. When orally administered to SNI-operated mice deficient in Nrf2 (*Nfe2l2*^-/-^), **1c** failed to reverse mechanical allodynia, compared to littermate wildtype controls (Fig. 2f, *P* < 0.001). There were no genotype differences in baseline thresholds or acute morphine analgesia (Fig. S4g,h; *P* = 0.597 (mechanical allodynia); *P* = 0.584 (dynamic allodynia)), excluding genotype as a confounding variable. These data indicate that Nrf2 mediates the antinociceptive effects of compound **1c**.

Unilateral peripheral nerve injury induces oxidative/nitrative stress in the ipsilateral sciatic nerve and L4/5 dorsal root ganglia (DRG; Fig. 2a)^10,11,27^. We tested whether **1c** would exclusively activate Nrf2 in these tissues as an indicator of selective cleavage of MMF in response to local oxidative/nitrative stress. Compound **1c** induced Nrf2 nuclear translocation (activation) only in the sciatic nerve injury site (Fig. S5a, *P* < 0.001) and ipsilateral DRG (Fig. 2g, *P* < 0.001), accompanied by selectively increased expression of Nrf2 target antioxidant genes in the ipsilateral DRG (Fig. 2h). In contrast, diroximel fumarate non-selectively activated Nrf2 and increased expression of antioxidant genes in the contralateral tissues as well (Fig. 2g, Fig. S5a, *P* < 0.001). Further demonstrating site-selective activation in contrast to diroximel fumarate, compound **1c** did not induce Nrf2 nuclear translocation at other unwanted sites (Fig. S5b; liver (*P* = 0.663), kidney (*P* = 0.909), lung (*P* = 0.999). Compound **5b** did not increase nuclear levels of Nrf2 in ipsilateral DRG, confirming that MMF generation by **1c** was responsible for Nrf2 translocation (Fig. S5c; *P* = 0.999).

Leukopenia is a common unwanted effect related to the systemic distribution of MMF^4,28^. We assessed whether administration of compound **1c** would have reduced immunosuppressive consequences as further evidence of spatially restricted pharmacodynamics. Diroximel fumarate reduced the total leukocyte count, as expected (Fig. 2i; *P* = 0.039). However, oral treatment with **1c** did not reduce total leukocyte count, compared to mice treated with vehicle (*P* = 0.651), indicating reduced immunosuppression.

## DISCUSSION

We have leveraged the Baeyer-Villiger chemical reaction in a biological setting to selectively deliver a small molecule Nrf2 activator in response to localized oxidative stress. Masking MMF with a 1,2-dicarbonyl moiety to allow generation at sites of pathology addresses the long-standing issue of side effects caused by systemic distribution of electrophilic drugs^2,3^. Importantly, 1,2-dicarbonyl compound **1c** is only cleaved by pathological concentrations of hydrogen peroxide or peroxynitrite, which is expected to leave physiological redox signalling intact^29^. The selective cleavage activates Nrf2 at sites of pathology *in vivo*, which mediates anti-nociceptive efficacy. Compound **1c** could also exert therapeutic effects by scavenging peroxides when participating in Baeyer-Villiger oxidation.

Compound **1c** and other 1,2-dicarbonyl compounds are suitable drug candidates, as they have broad synthetic scope to optimize pharmacokinetic and toxicological properties. The diversity offers further advantage in that the cleaved carboxylic, carbonic, or carbamic acid fragment **3** (Fig. 1a) can be relatively low in molecular weight and selected to be biologically benign and non-toxic.

We show that oral delivery of compound **1c** to activate Nrf2 at local sites of oxidative stress reverses chronic pain of diverse aetiologies in preclinical models, consistent with Nrf2-dependent antinociceptive effects of fumarates^26^. Nrf2 activation is an attractive therapeutic target because its activation increases levels of the panoply of endogenous antioxidants that scavenge free radicals and their precursors, which underlie chronic pain pathology^10,11,27^. Activating Nrf2 with a single small molecule is therefore superior to supplementation of individual antioxidants that generally have unfavorable pharmacokinetic and pharmacodynamics profiles^3,10,11^. Chronic pain remains a large unmet medical need, and non-addictive treatments that engage endogenous pain resolution mechanisms like Nrf2 are a unique approach^30^.

Oxidative stress is a pathological process common to numerous diseases and disorders^1-3^. 1,2-Dicarbonyl **1c** therefore has potential to treat a wide range of pathologies. The 1,2-dicarbonyl platform could further have broad application in targeting other molecular payloads to sites of oxidative stress.

## METHODS

### 1,2-Dicarbonyl compound synthesis and characterisation

All anhydrous solvents were commercially obtained and stored in Sure-Seal bottles under nitrogen or transferred to an Inert Corporation Solvent Purification System and dispensed from there. All other reagents and solvents were purchased as the highest grade required and used without further purification. All organic extracts were dried over anhydrous magnesium sulfate (MgSO_4_). Thin-layer chromatography (TLC) used aluminum sheets coated with silica gel 60 F_254_ from Merck and were visualized using ultraviolet light. Melting points were taken on a Reichert Thermovar Kofler apparatus and are uncorrected. Infrared spectra were recorded on a Perkin Elmer Spectrum 400 FT-IR / FT-FIR Spectrometer as neat samples unless otherwise stated. ^1^H NMR and ^13^C NMR, spectra were acquired on an Agilent 500 MHz spectrometer or Agilent 600 MHz spectrometer. High resolution mass spectrometry (HRMS) was performed on an Agilent 6230 ESI-TOF LCMS. Analytical RP HPLC spectra were acquired on an Agilent 1100 Infinity series machine using a Phenomenex Luna C18(2) column (250 × 4.6 mm, 100Å, 5 μm) with a buffer system of 0.1% TFA in water (buffer A) and 0.1% TFA in acetonitrile (buffer B), elution was with a gradient of 0–100% A-B over 20 min. All yields reported refer to isolated material judged to be homogeneous by TLC, HPLC and NMR spectroscopy.

α-Keto amide **1b** was synthesized as reported^31^ and α-keto ester **1c** was synthesized from dimethyl 2-oxoglutarate **5b**, using the similar bromination / elimination conditions described for **1b** (Fig. S1a). Both compounds were stored as solids at 4°C in the absence of light to avoid UV light assisted isomerization.

We designed a synthetic approach to aromatic 1,2-diketones of type **1a** (Fig. S1b). *trans*-Cinnamaldehyde **6** reacted with stabilized phosphorus ylide **7** to give *trans, trans* 1,3-butadiene **8** as the major product. Purified diene **8** was then selectively dihydroxylated employing standard Sharpless conditions outlined for closely related compounds^32^. 1,2-Diketone **1a** was generated by treatment of purified diol **10** with Dess-Martin periodinane, neat. This technique was used to minimize competing oxidative cleavage of the diol to benzaldehyde and methyl (2*E*)-4-oxo-2-butenoate. 1,2-Diketone **1a** was found to isomerise to the *cis* isomer in deuterated chloroform on exposure to ambient light. Once solvent was removed and **1a** was stored without solvent it reverted to and remained as the *trans* isomer. 1,2-Diketone **1a** was fully characterized and its identity confirmed by subsequent reaction with hydrogen peroxide to generate benzoic acid and MMF. The IR and ^1^H NMR spectra we obtained did not match those reported in the literature (Tables S3), indicating **1a** was not the product obtained from the high-pressure palladium catalysed double carbonylative vinylation of 4-iodobenzene and methyl acrylate in the presence of carbon monoxide, as reported^33^. Ours is therefore the first reported synthesis of 1,2-diketones of type **1a** (Fig. S1b).

α-Keto acid **1d** was synthesised from **1c** obtaining it pure by extraction from its hydrolysis reaction in 100 mM phosphate buffer (pH7.4).

### In vitro Nrf2 reporter assay

NRF2/ARE luciferase reporter HEK293 cells (SL-0042-NP, Signosis, Santa Clara, USA) were maintained in Dulbecco’s Modified Eagle’s Medium (DMEM) (SH30243.01, Cytiva, Marlborough, USA) supplemented with 10% Fetal Bovine Serum (FBS) (03-600-511, ThermoFisher Scientific, Waltham, USA) and 1% Penicillin/Streptomycin (SV30010, Cytiva). Cells were seeded into 48-well flat bottom cell culture plate (3548, Corning, Corning, USA) in 500 μL of supplemented DMEM at a concentration of 2 × 10^5^ cells/mL and incubated overnight at 37°C with 5% CO_2_ in a humidified environment. Prior to treatment, the media was replaced with fresh 0.1% FBS in DMEM. Tert-Butylhydroquinone (tBHQ; 0, 0.5-30 μM; 112941, Sigma-Aldrich, St Louis, USA), a well-known Nrf2 activator, was used as a positive control to confirm Nrf2-dependent luciferase production (Fig. S6). To identify concentrations of hydrogen peroxide and peroxynitrite that did not activate Nrf2 *per se*, cells were first treated with concentration ranges of hydrogen peroxide (H1009, Sigma-Aldrich) or peroxynitrite (20-107, Sigma-Aldrich) (0, 1-300 μM) (Fig. S5). In the next series of experiments, cells were treated with concentration ranges of **1a** (synthesized by T. Avery), **1b** (synthesized by T. Avery), **1c** (synthesized by T. Avery), **5b** (349631, Sigma-Aldrich), or monomethyl fumarate (651419, Sigma-Aldrich) (0, 1-100 μM), followed by a fixed concentrations of hydrogen peroxide (1 or 10 μM), peroxynitrite (1 or 20 μM), or media control. These concentrations respectively represent physiological and pathological levels^15,16^. Cells were incubated with the treatments for 16 hours. After washing with phosphate buffer saline (PBS) (ThermoFisher Scientific), cells were lysed by a 15-min incubation at room temperature with passive lysis buffer (E1941, Promega, Madison, USA). Cell lysates (30 μL) were transferred to 96-well white/clear flat bottom plate (3632, Corning), and mixed with 150 μL of luciferase substrate (LUC100, Signosis). The plates were read in a Synergy HTX Multi-Mode Reader (BioTek, Winooski, USA). All conditions were performed in triplicate.

### Animals

Pathogen-free adult male and female C57BL/6J mice (8 weeks old on arrival; The Jackson Laboratory, Bar Harbor, USA) were used. Male and female (8 to 12 weeks old) wild type and *Nfe2l2*^-/-^ mice on a C57BL/6J genetic background (The Jackson Laboratory) were bred at The University of Texas MD Anderson Cancer Center (Houston, USA). Mice were housed five per cage in a light- and temperature-controlled room (12:12-h light-dark cycle, light on at 7:00 AM) with food and water available *ad libitum*. All procedures were approved by the MD Anderson Cancer Center Animal Care and Use Committee.

### Spared nerve injury (SNI) surgery

SNI^18^ was performed as described for mice^34^. Briefly, under inhaled isoflurane anesthesia, the tibial and common peroneal nerves were isolated, tightly ligated with 6-0 silk (707G, Ethicon, Somerville, USA), and transected immediately distal to the ligation. The sural nerve was left intact. For sham surgery, the nerves were exposed, but not ligated or transected. Animals were monitored post-operatively until fully ambulatory prior to return to their home cage.

### Surgical destabilization of the medial meniscus (DMM)

Mice were subjected to the DMM model of osteoarthritis in one hindlimb, as previously described^21^. Mice were anesthetized with isoflurane and placed in dorsal recumbency. Carprofen (5mg/kg, SQ) was administered immediately prior to beginning surgery. A 3 mm longitudinal incision was made over the right patella, and the joint capsule was opened with micro Vannas scissors under 4X magnification. Blunt dissection of the infrapatellar fat pad exposed the meniscotibial ligament of the medial meniscus, which was then transected using a #11 scalpel blade. The incision was closed with two 9mm AutoClip wound clips, which were removed 10 days post-surgery. Sham surgery consisted of the same anesthesia, analgesia and skin/joint capsule incisions, without transection of the meniscotibial ligament. Animals were monitored post-operatively until fully ambulatory prior to return to their home cage.

### In vivo drug treatments

Cisplatin (TEVA Pharmaceuticals, North Wales, PA) was diluted in sterile saline and administered for 5 days (2.3 mg/kg/day, i.p.) followed by 5 days of rest and a second round of 5 doses to induce chemotherapy-induced neuropathy (CIPN)^19^. Compound **1c** (MW= 172.14) and diroximel fumarate (MW= 255.23) (synthesized by T. Avery) were suspended in methylcellulose (viscosity 15 cP, 2% w/v in water; Sigma-Aldrich). For SNI anti-nociception: **1c** (100, 225, or 350 μmol/kg/day, p.o.), **1d** (350 μmol/kg/day, p.o.), diroximel fumarate (350 μmol/kg/day, p.o.) or methylcellulose or phosphate buffer vehicle controls (equivolume, p.o.) were administered daily, beginning 7 days after SNI/sham, and continuing for 3 (reflex tests) or 7 days (conditioned place preference tests). For CIPN anti-nociception: **1c** (350 μmol/kg/day, p.o.) or methylcellulose vehicle were administered for 5 consecutive days, beginning 3 days after the last cisplatin dose. For anti-nociceptive tolerance: **1c** (350 μmol/kg/day, p.o.) or the positive control morphine sulfate (6 mg/kg/day, s.c.; gifted from the National Institute on Drug Abuse Drug Supply Program, Research Triangle Institute, NC) were administered daily for 5 days, beginning 7 days after SNI. For acute morphine analgesia: morphine sulfate (5 mg/kg, s.c.; gifted from the National Institute on Drug Abuse Drug Supply Program). For leukopenia assessment: naïve mice were treated with **1c** (350 μmol/kg/day, p.o.), diroximel fumarate (350 μmol/kg/day, p.o.), or methylcellulose vehicle were administered daily for 10 days.

### Behavioural sensory testing

All behavioural tests were conducted by an experimenter who was blinded to group assignments. Mice received at least three 60-min habituations to the test environment before reflex testing. Rodents were placed in a small plexiglass enclosure on a mesh stand. Tactile allodynia was measured using the von Frey test as described previously^26^. The 50% paw withdrawal threshold was determined using the “up-down” method^35^.

Dynamic allodynia was measured by lightly stroking the plantar surface of the hindpaw with a soft paintbrush, using a protocol and scoring system developed by Dr. Enrique José Cobos (personal communication), as previously reported^36^. A paintbrush (5/0, Princeton Art & Brush Co.) was prepared by blunting the tip and removing the outer layer of hairs. The lateral plantar region of the left hindpaw (sural nerve territory) was stimulated by light stroking (∽2 cm/s) with the paintbrush, in the direction from heel to toe. The paw withdrawal response was scored according to the following criteria, score = 0: walking away or occasionally very brief paw lifting (≤ 1 s); score = 1: sustained lifting (> 2 s) of the stimulated paw toward the body; score = 2: a strong lateral lifting above the level of the body; score = 3: flinching or licking of the affected paw. Average scores for each mouse were obtained from three stimulations at intervals of at least 3 min.

Spontaneous pain was tested using a conditioning paradigm with retigabine (#R-100; Alomone Laboratory, Jerusalem, Israel) as the conditioned stimulus to briefly relieve pain, as previously described^19,25,37^. Mice were first allowed to freely explore the conditioned place preference apparatus, consisting of two chambers (one dark, one light) connected by a hallway (Stoelting, Wood Dale, USA), for 15 minutes. The time spent in the light chamber was recorded. During the conditioning phase, mice were first administered saline (i.p.) and kept in the dark chamber for 20 minutes. Three hours later, the analgesic retigabine was administered (10 mg/kg; i.p.) and after 10 mins, the mice were placed in the light chamber for 20 minutes. The conditioning was completed over four consecutive days. On the fifth day, the mice were again allowed to freely explore the apparatus for 15 minutes without any retigabine/saline injections. Data are presented as the difference in time spent in the light (retigabine-paired) chamber during the drug-free test on day five minus time spent in the light chamber at baseline (pre-conditioning phase). A mouse with spontaneous pain should show an increase in time spent in the light chamber that was paired with retigabine than it did in the preconditioning phase.

To examine numbness, we used a modified protocol of the adhesive removal test, as described^19,38^. Briefly, a round adhesive patch (3/16” Teeny Tough-Spots; USA Scientific INC, Ocala, USA) was placed on the plantar surface of the hind paws. The latency to attend to the patch (e.g., shaking or attempted removal) was recorded within a 15-minute testing time.

### Tissue collection

Within 4□h of the final dose, mice were deeply anesthetized with Beuthanasia-D (Merck, Kenilworth, USA) and then transcardially perfused with ice-cold saline. In some experiments, blood was collected via cardiac puncture prior to perfusion. The ipsilateral and contralateral sciatic nerve (5 mm proximal to the transection), L4/5 dorsal root ganglia (DRG), liver, kidney, and lung were isolated and rapidly frozen for subsequent analysis.

### qPCR

Total RNAs were extracted from HEK293 cells or DRG tissues using TRIzol (ThermoFisher Scientific). One μg RNA was used for reverse transcription with iScript Reverse Transcription Supermix (Bio-Rad). Real-time polymerase chain reaction was carried out in a final volume of 20 μl with iTaq Universal SYBR Green Supermix (Bio-Rad) containing 2 μl of five times diluted cDNA and monitored by CFX Connect Real-Time PCR Detection System (Bio-Rad). The following cycling parameters were used: 95°C for 3 min, 40 cycles of 95°C for 5 s, and 60°C for 30 s. Primer sequences are reported in Table S7. The level of the target mRNA was quantified relative to the house-keeping gene (*Gapdh*) using the ΔΔCT method. *Gapdh* was not significantly different between treatments.

### Western blotting

DRG from 3 mice or sciatic nerves from two mice were pooled within groups (treatment, lateralization, sex) to ensure that sufficient protein could be obtained for analysis. Liver, kidney, and lung were not pooled. Nuclear fractions were isolated with a NE-PER Nuclear and Cytoplasmic Extraction Kit (78835, ThermoFisher Scientific), according to manufacturer instructions. Western blotting was performed as previously described^3^. Nuclear proteins were subjected to NuPAGE Bis-Tris (4 to 12%) gel electrophoresis under reducing conditions. After transfer to nitrocellulose membranes (IB23001, Invitrogen, USA), non-specific binding sites were blocked with Superblock buffer (37515, ThermoFisher Scientific) for 1 hour at room temperature. Membranes were incubated overnight at 4°C with primary anti-Nrf2 antibody (1:1000; rabbit polyclonal IgG; ab31163, Abcam) and anti-histone H3 antibody (1:2000; rabbit polyclonal IgG; ab1791, Abcam) (loading control). The membranes were then washed with phosphate buffered saline containing 0.1% Tween-20 and probed with horseradish peroxidase secondary antibody (1:5,000; goat polyclonal IgG; Jackson ImmunoResearch, West Grove, USA) in blocking buffer containing 0.1% Tween-20 for 1□h at room temperature. After washing with 1 x PBS containing 0.1% Tween-20, membranes were developed with enhanced chemiluminescent substrate (ThermoFisher Scientific). Images were acquired using ImageQuant LAS 4000 (GE Healthcare Life Sciences, USA).

Densitometry analysis was performed using ImageQuant TL software (GE Healthcare Life Sciences). Data were normalized to loading control (histone H3).

### Leukocyte counting

Leukocytes from cardiac blood were stained with Türk’s solution (Sigma-Aldrich) according to manufacturer instructions, and manually counted on a hemocytometer by an experimenter who was blinded to treatment conditions.

### Statistics

Differences between *in vitro* concentration-response relationships were determined by comparing the slopes of fitted functions for shared parameters. Linear models were selected as the pharmacologically-relevant concentrations tested were within the linear range of the concentration-response functions. Von Frey and brush data were analysed by repeated measures two- or three-way ANOVA with Tukey *post hoc* tests where appropriate. Conditioned place preference data were analysed by two-way ANOVA with Tukey *post hoc* tests. Data from the adhesive removal test were analysed by unpaired T test. Gene expression, protein levels, and leukocyte numbers were analysed by one-way ANOVA followed by Dunnett or Tukey *post hoc* tests as appropriate. Analyses were performed using Prism 9 (GraphPad). *P* < 0.05 was considered statistically significant. Data are expressed as mean ± SD.

## ACKNOWLEDGEMENTS

This work is supported by National Institutes of Health grants RF1 NS113840 (P.M.G.), and in part by P30 CA016672 (MD Anderson Cancer Center); and, Australian Research Council grants CE140100003 and DP180101581 (A.D.A.).

## AUTHOR CONTRIBUTIONS

T.D.A., P.M.G., and A.D.A. conceived the study. T.D.A, J.L., D.J.L.T., P.M.G., and A.D.A. designed most of the experiments, and A.J.S. and J.Y. designed certain experiments. T.D.A., J.L., D.J.L.T., F.R.C., M.S.U.R., C.A, J.Y. collected the data. T.D.A, P.M.G., and A.D.A. drafted the paper. All authors contributed to data analysis and writing of the paper.

---

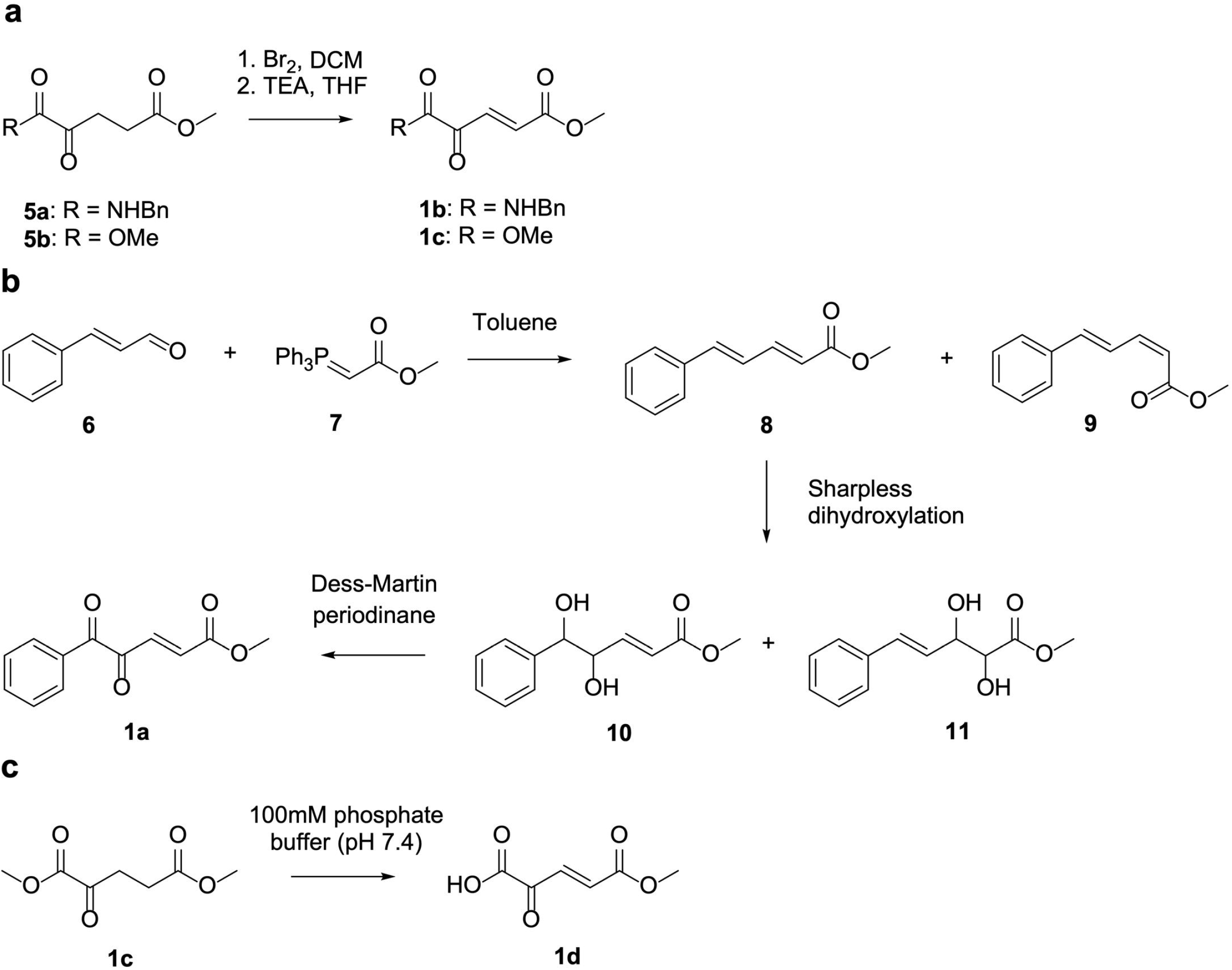

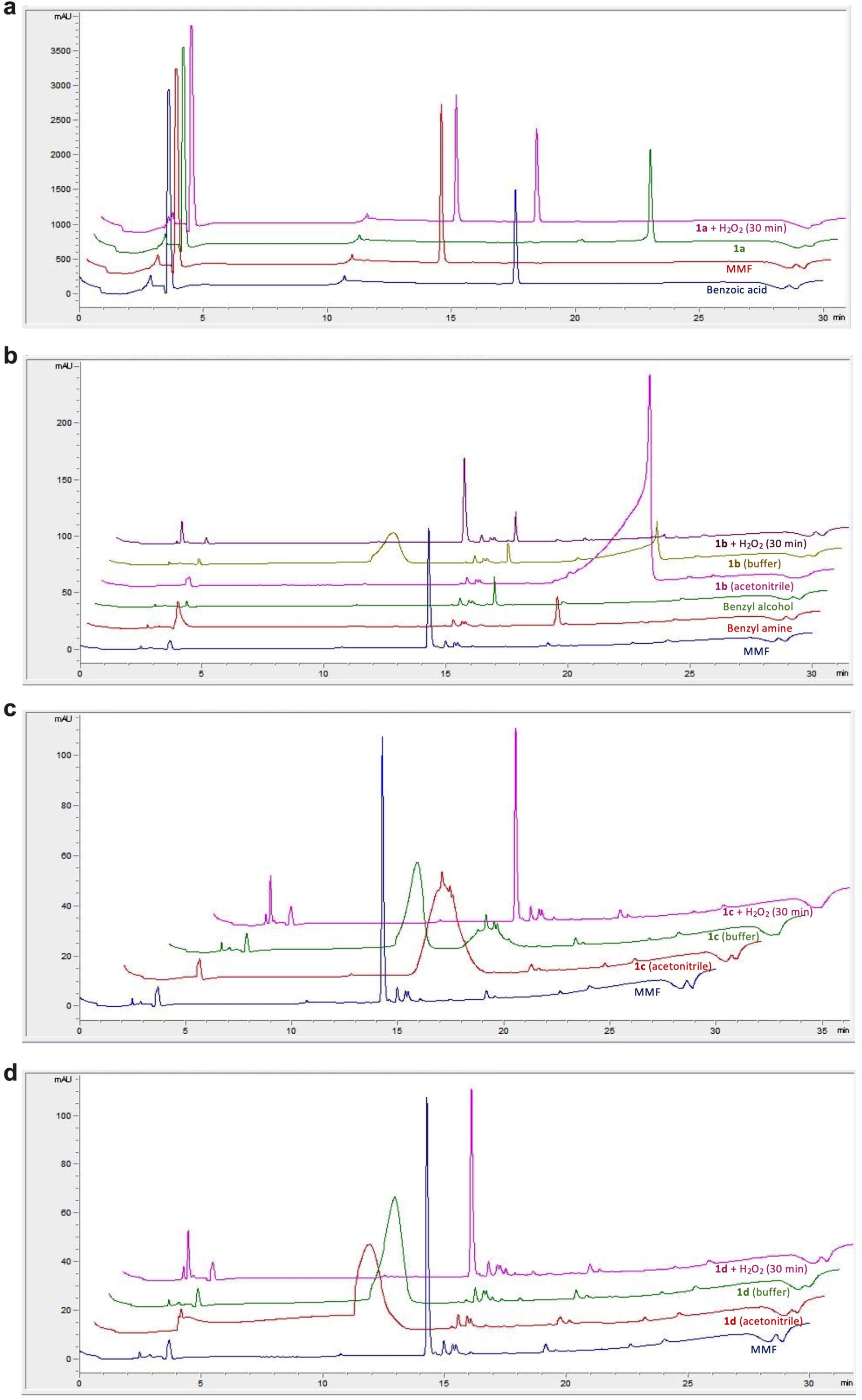

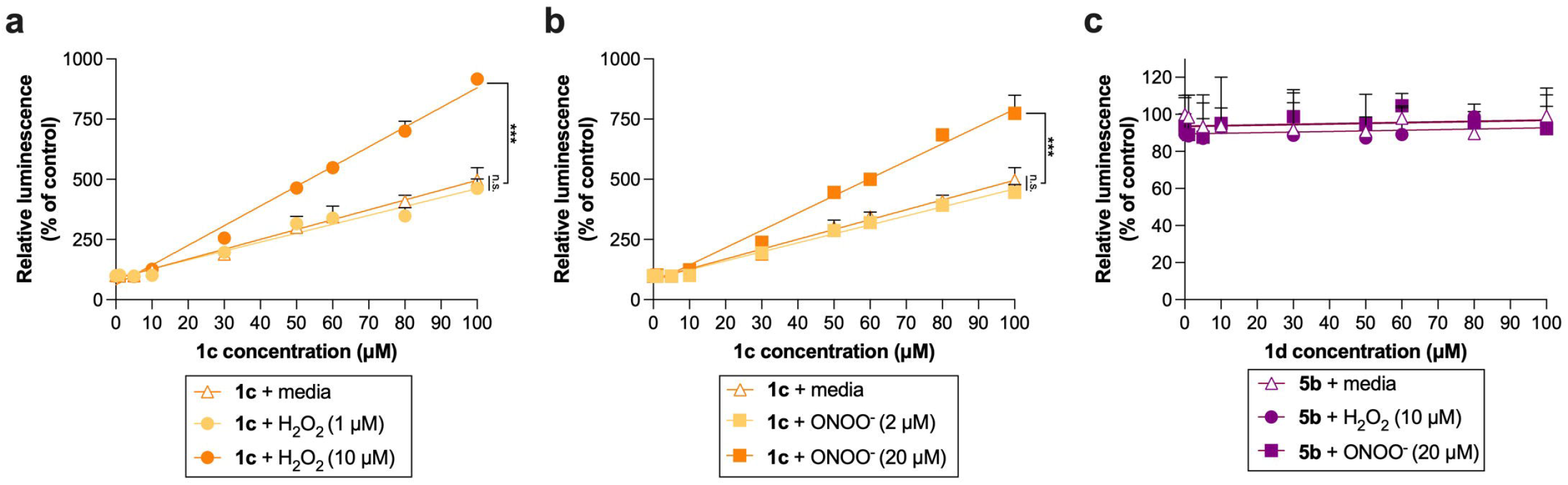

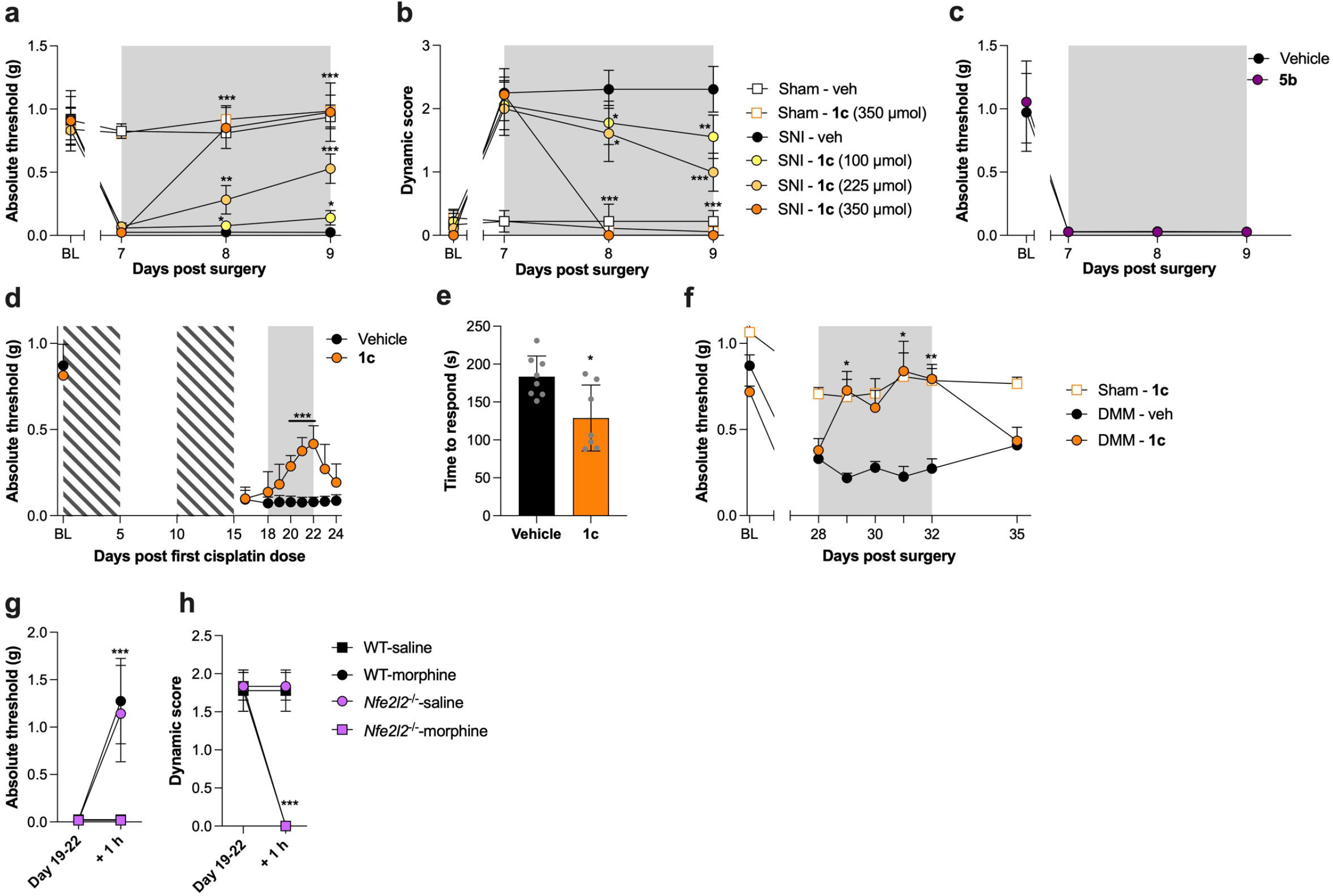

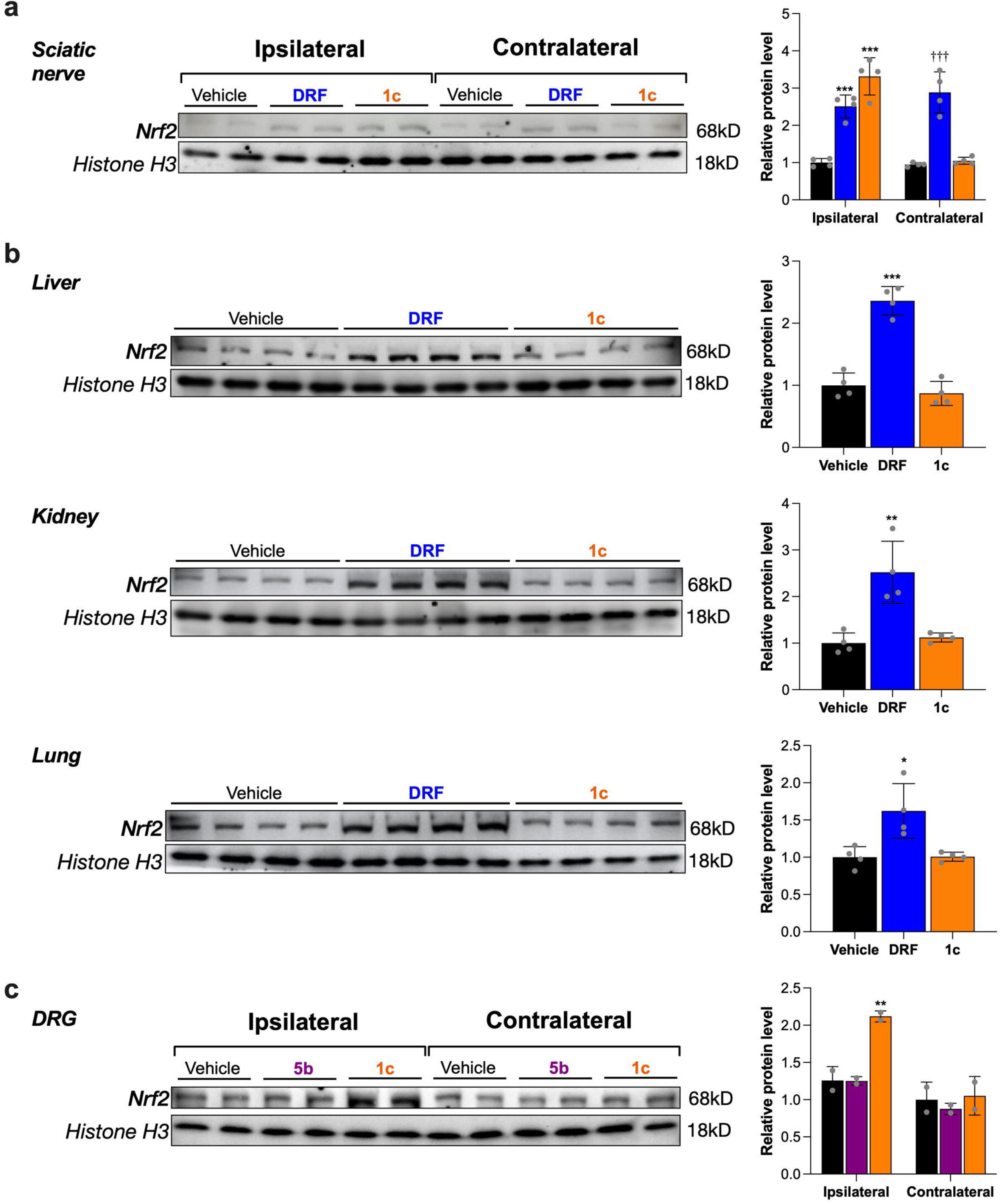

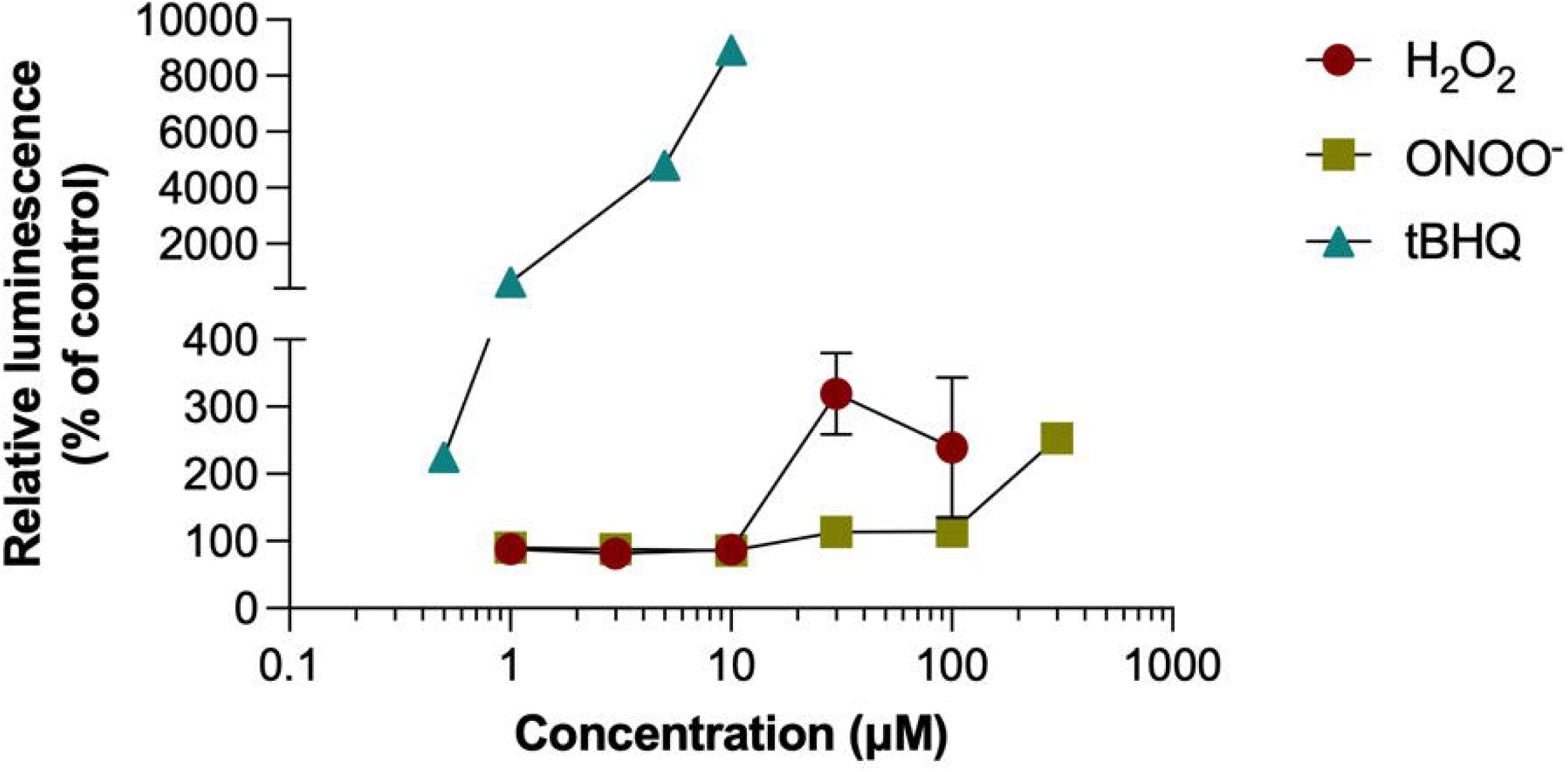

