## Supplementary Figures and Tables for "Drug targeting to sites of oxidative stress using the Baeyer-Villiger reaction"

**Figure S1.** Synthetic routes to 1,2-dicarbonyl compounds **1a-d**. In case the structures contain one or more stereogenic centres, the respective structure is depicted in an arbitrary configuration. There structures depict single enantiomers as well as mixtures of enantiomers in all ratios and /or mixtures of diastereomers in all ratios.

**Figure S2.** Stacked HPLC traces of authentic samples and reaction of **1a** (0.33 minute offset), **1b** (0.33 minute offset), **1c** (2 minute offset), or **1d** (0.67 minute offset) with hydrogen peroxide in buffer. (**a**) Authentic samples of MMF, benzoic acid and **1a** were made up as 30 mM stock solutions in DMSO and diluted to give 1 mM solutions in 100 mM phosphate buffer (pH 7.4). All samples were run individually on the HPLC to reference the reaction mixture. 33 μl of **1a** DMSO stock was added to 967 μl of 100 mM phosphate buffer (pH 7.4) containing hydrogen peroxide at a concentration of 5.15 mM. The resulting 1 mM solution of **1a** with a 5x excess of hydrogen peroxide was then run on the HPLC 30 minutes after addition of **1a** to the hydrogen peroxide in phosphate buffer and visualized at 220 nm. (**b**) Authentic samples of MMF, benzyl amine, benzyl alcohol and **1b** were made up as 30 mM stock solutions in DMSO and diluted to give 1 mM solutions in 100 mM phosphate buffer (pH 7.4). Another 1 mM sample of **1b** was made up in acetonitrile. All samples were run individually on the HPLC to reference the reaction mixture. 33 μl of **1b** DMSO stock was added to 967 μl of 100 mM phosphate buffer (pH 7.4) containing hydrogen peroxide at a concentration of 5.15 mM. The resulting 1 mM solution of **1b** with a 5 x excess of hydrogen peroxide was then run on the HPLC 30 minutes after addition of **1b** to the hydrogen peroxide in phosphate buffer and visualised at 254 nm. (**c**) Authentic samples of MMF and **1c** were made up as 30 mM stock solutions in DMSO and diluted to give 1 mM solutions in 100 mM phosphate buffer (pH 7.4). Another 1 mM sample of **1c** was made up in acetonitrile. All samples were run individually on the HPLC to reference the reaction mixture. 33 μl of **1c** DMSO stock was added to 967 μl of 100 mM phosphate buffer (pH 7.4) containing hydrogen peroxide at a concentration of 5.15 mM. The resulting 1 mM solution of **1c** with a 5 x excess of hydrogen peroxide was then run on the HPLC 30 minutes after addition of **1c** to the hydrogen peroxide in phosphate buffer and visualised at 254 nm. (**d**) Authentic samples of MMF and **1d** were made up as 30 mM stock solutions in DMSO and diluted to give 1 mM solutions in 100 mM phosphate buffer (pH 7.4). Another 1 mM sample of **1d** was made up in acetonitrile. All samples were run individually on the HPLC to reference the reaction mixture. 33 μl of **1d** DMSO stock was added to 967 μl of 100 mM phosphate buffer (pH 7.4) containing hydrogen peroxide at a concentration of 5.15 mM. The resulting 1 mM solution of **1d** with a 5 x excess of hydrogen peroxide was then run on the HPLC 30 minutes after addition of **1d** to the hydrogen peroxide in phosphate buffer and visualised at 254 nm.

**Figure S3. *In vitro* characterization of 1,2-dicarbonyl compound 1c with physiological concentrations of peroxides.** NRF2/ARE luciferase reporter HEK293 cells were treated with **(a-b)** compound **1c** together with media, **(a)** H_2_O_2_ (1, 10 μM), or (**b**) ONOO^-^ (2, 20 μM); or **(c)** compound **5b** together with media, H_2_O_2_ (10 μM), or ONOO^-^ (20 μM). ARE-luciferase activity quantified; not significant (n.s.), ****P* < 0.001.

**Figure S4. *In vivo* assessment of 1,2-dicarbonyl prodrug efficacy.** **(a-c)** Beginning 7 days after SNI or sham surgery, male (n=3) and female (n=3) mice were treated with **(a-b)** oral **1c** (100, 225, or 350 μmol/kg/day), **(c)** oral **5b** (350 μmol/kg/day), or vehicle every day for 3 days (gray box). **(a, c)** Mechanical allodynia and **(b)** dynamic allodynia were assessed. Relative to vehicle: **P* <0.05, ***P* <0.01, ****P* <0.001. **(d-e)** Cisplatin was administered for 5 days (2.3 mg/kg/day, i.p.) followed by 5 days of rest and a second round of 5 doses to induce chemotherapy-induced neuropathy (hatched boxes). **(d)** Male (n=3) and female (n=3) mice were treated with oral **1c** (350 μmol/kg/day) for 5 consecutive days, beginning 3 days after the last cisplatin dose. Mechanical allodynia was assessed over a timecourse, and **(e)** the adhesive removal test for numbness was performed 4 h after the last ***1c*** dose. Relative to vehicle: **P* <0.05; ****P* <0.001. **(f)** Beginning 28 days after DMM or sham surgery, male (n=4) and female (n=4) mice were treated with oral **1c** (350 μmol/kg/day) or vehicle every day for 4 days (gray box). Mechanical allodynia was assessed. 1c vs. vehicle: **P* <0.05, ***P* <0.01. **(g-h)** After other drug treatments had washed out on days 19-22 after SNI, male and female *Nfe2l2*^-/-^ and wildtype control mice (n=3/sex/group) were treated with morphine (5 mg/kg), and **(g)** mechanical allodynia and **(h)** dynamic allodynia were assessed 1 h later. Relative to vehicle controls: ****P* <0.001.

**Figure S5. Post-mortem assessment of Nrf2 nuclear translocation.** **(a-c)** Beginning 7 days after SNI surgery, mice were treated with oral diroximel fumarate (350 μmol/kg/day), or **1c** (350 μmol/kg/day), **5b** (350 μmol/kg/day), or vehicle every day for 3 days. **(a)** Sciatic nerves from 2 mice were pooled (within sexes), and nuclear extracts were probed for Nrf2 (n = 2/sex per group). **(b)** Nuclear extracts from liver, kidney, and lung were probed for Nrf2 (n = 2/sex per group). **(c)** L4/5 DRG from 3 mice were pooled after 3 days of treatment, and nuclear extracts were probed for Nrf2 (n = 2/sex per group). Quantification by densitometry is presented for each blot. Relative to ipsilateral vehicle: **P* <0.05, ***P* <0.01, ****P* <0.001; relative to contralateral vehicle and **1c**: ^†††^*P* <0.001.

**Figure S6. *In vitro* optimization of peroxide concentrations.** NRF2/ARE luciferase reporter HEK293 cells were treated with concentration ranges of H_2_O_2_ or ONOO^-^ to determine the concentrations at which peroxides would activate Nrf2 *per se*. Cells were treated with tBHQ as a positive control. ARE-luciferase activity was quantified.

**SUPPLEMENTARY TABLES**

**Table S1.** Hydration equilibria for 1,2-dicarbonyl compounds **1a-d** and **5b** at 22°C determined by ^1^H NMR.

| 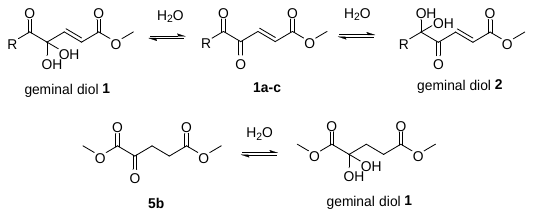 | | | |
| --- | --- | --- | --- |
| **Compound** | **Relative ratio** | | |
|  | **1,2-Dicarbonyl form** | **Geminal diol 1** | **Geminal diol 2** |
| **1a^a^** | 1 | 0.07 | 0.04 |
| **1a^b^** | 1 | 0.6 | 0.4 |
| **1b^a^** | 1 | 0.79 | - |
| **1c^a^** | 1 | 0.84 | - |
| **1c^c^** | 1 | 5.9 | - |
| **1d^c^** | 1 | 2 | - |
| **5b^c^** | 1 | 2 | - |

^a^Spectra recorded in 2:1 acetonitrile-*d*_3_ : D_2_O. ^b^Spectra recorded in 2:1 DMSO-*d*_6_ : D_2_O ^c^Spectra recorded in D_2_O.

**Table S2.** Equilibria established between 1,2-dicarbonyl compounds **1a-d** and **5b,** and their geminal diols and Criegee intermediates upon reaction with hydrogen peroxide at 22°C determined by ^1^H NMR.

| 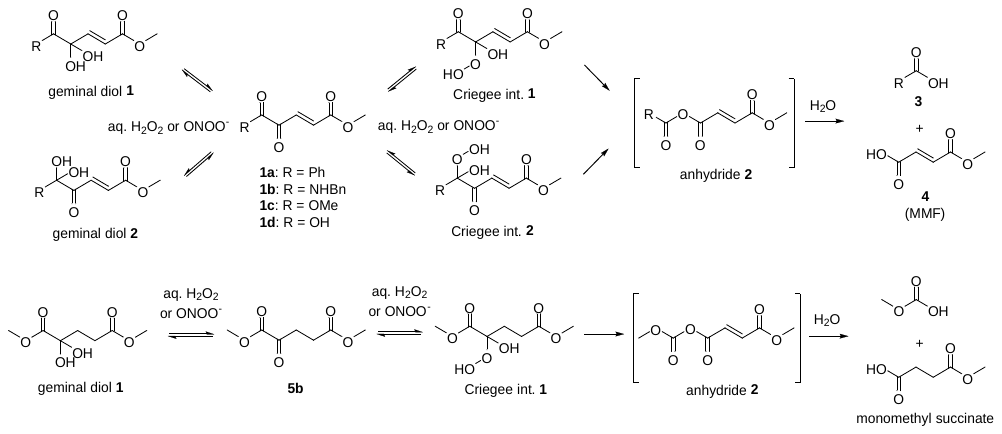 | | | | | |
| --- | --- | --- | --- | --- | --- |
| **Compound** | **Relative ratio** | | | | |
|  | **1,2-Dicarbonyl form** | **Geminal diol 1** | **Geminal diol 2** | **Criegee Intermediate 1** | **Criegee Intermediate 2** |
| **1a^a^** | 1 | 0.11 | 0.11 | 0.52 | 0.54 |
| **1b^a^** | 1 | 0.79 | - | 4.71 | - |
| **1c^a^** | 1 | 0.82 | - | 5.25 | - |
| **1c^b^** | 1 | 5.88 | - | 10.74 | - |
| **1d^b^** | 0 | 1 | - | Reaction too rapid to observe Criegee intermediate | - |
| **5b^b^** | 1 | 2 | - | 4.4 | - |

^a^Spectra recorded in 2:1 acetonitrile-*d*_3_ : D_2_O. ^b^Spectra recorded in D_2_O.

**Table S3.** ^1^H and ^13^C NMR of 1,2-dicarbonyl and geminal diols of **1a-d** and **5b** at 22°C.

| 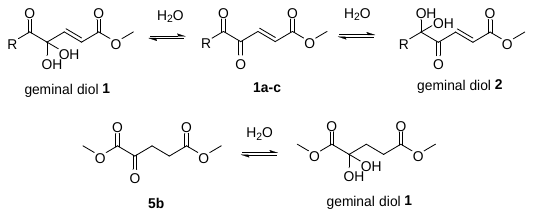 | | | | | | |
| --- | --- | --- | --- | --- | --- | --- |
| **Compound** | **1,2-Dicarbonyl form** | | **Geminal diol 1** | | **Geminal diol 2** | |
|  | **^1^H NMR (500 MHz)** | **^13^C NMR**  **(125 MHz)** | **^1^H NMR (500 MHz)** | **^13^C NMR (125 MHz)** | **^1^H NMR (500 MHz)** | **^13^C NMR (125 MHz)** |
| **1a^a^** | δ 8.00 (d, *J* = 8.0 Hz, 1H), 7.68 (t, *J* = 7.5 Hz, 1H), 7.54-7.49 (m, 3H), 6.90 (d, *J* = 16.0 Hz, 1H), 3.85 (s, 3H) | δ 191.0, 190.9, 165.2, 135.4, 135.3, 135.1, 132.1, 130.3, 129.0, 52.6 | - | - | - | - |
| **1a^b^** | δ 8.02 – 7.96 (m, 2H), 7.79 – 7.72 (m, 1H), 7.63 – 7.56 (m, 2H), 7.52 (d, *J* = 16.1 Hz, 1H), 6.83 (d, *J* = 16.1 Hz, 1H), 3.78 (s, 3H). | δ 190.18, 189.50, 165.02, 135.64, 134.76, 133.16, 131.86, 130.25, 128.83, 52.41 | - | - | - | - |
| **1a^c^** | δ 7.95–7.90 (m, 2H), 7.74–7.67 (m, 1H), 7.57–7.50 (m, 2H), 7.40 (d, *J* = 16.0 Hz, 1H), 6.82 (d, *J* = 16.1 Hz, 1H), 3.75 (s, 3H) | δ 191.77, 191.64, 165.90, 135.44, 135.32, 135.15, 131.89, 130.24, 129.17, 52.62. | Olefinic peaks δ 6.97 (d, *J* = 15.7 Hz, 1H), 6.25 (d, *J* = 15.7 Hz, 1H) | - | Olefinic peaks δ 7.29 (d, *J* = 15.8 Hz, 1H), 6.70 (d, *J* = 15.8 Hz, 1H) | - |
| **1a^d^** | δ 8.00 – 7.96 (m, 2H), 7.82 – 7.77 (m, 1H), 7.65-7.61 (m, 2H), 7.48 (d, *J* = 16.2 Hz, 1H), 6.86 (d, *J* = 16.2 Hz, 1H), 3.80 (s, 3H) | δ 192.40, 191.98, 166.64, 136.51, 136.26, 135.67, 132.52, 131.26, 130.23, 53.79 | δ 8.18 – 8.14 (m, 2H), 7.70 – 7.65 (m, 1H), 7.58 – 7.52 (m, 2H), 7.03 (d, *J* = 15.7 Hz, 1H), 6.25 (d, *J* = 15.7 Hz, 1H), 3.69 (s, 3H) | δ 197.24, 167.58, 148.45, 134.94, 134.03, 131.34, 129.66, 123.27, 95.75, 53.11 | δ 7.57 – 7.52 (m, 2H), 7.46 – 7.41 (m, 3H), 7.37 (d, *J* = 15.9 Hz, 1H), 6.70 (d, *J* = 15.9 Hz, 1H), 3.72 (s, 3H) | δ 196.64, 166.81, 140.43, 135.86, 132.57,130.19, 129.66, 127.43, 97.15, 53.57 |
| **1b^a^** | δ 7.97 (d, *J* = 16.0 Hz, 1H), 7.39 – 7.31 (m, 3H), 7.31 – 7.26 (m, 3H), 7.05 (d, *J* = 16.0 Hz, 1H), 4.67-4.37 (m, 2H), 3.84 (s, 3H) | δ 182.42, 165.24, 160.59, 135.56, 134.32, 134.29, 129.13, 128.94, 128.92, 68.61, 52.72. | - | - | - | - |
| **1b^c^** | δ 7.48 (d, *J* = 16.0 Hz, 1H), 7.43-7.29 (m, 5H), 6.84 (d, *J* = 16.0 Hz, 1H), 5.27 (s, 2H), 3.75 (m, 3H) | δ 182.76, 165.86, 160.35, 134.73, 134.55, 134.44, 128.91, 128.81, 128.71, 68.38, 52.59 | δ 7.43-7.29 (m, 5H), 6.84 (d, *J* = 16.0 Hz, 1H), 6.18 (d, *J* = 15.6 Hz, 1H), 5.16 (s, 2H), 3.67 (s, 2H) | δ 170.16, 167.08, 145.07, 135.40, 128.76, 128.57, 128.23, 122.62, 91.81, 67.74, 51.96 | - | - |
| **1c^a^** | δ 7.63 (d, *J* = 16.0 Hz, 1H), 6.98 (d, *J* = 16.0 Hz, 1H), 3.94 (s, 3H), 3.85 (s, 3H) | δ 182.30, 165.23, 161.10, 135.58, 134.24, 53.51, 52.75. | - | - | - | - |
| **1c^c^** | δ 7.50 (d, *J* = 16.0 Hz, 1H), 6.86 (d, *J* = 16.0 Hz, 1H), 3.84 (s, 3H), 3.76 (s, 3H), | δ 182.76, 165.96, 161.07, 134.52, 134.47, 53.31, 52.61 | δ 6.83 (d, *J* = 15.7 Hz, 1H), 6.19 (d, *J* = 15.8 Hz, 1H), 3.71 (s, 3H), 3.69 (s, 3H), | δ 170.92, 167.22, 145.07, 122.54, 91.72, 53.11, 52.01 | - | - |
| **1c^e^** | δ 7.67 (d, *J* = 16.0 Hz, 1H), 6.99 (d, *J* = 16.0 Hz, 1H), 3.94 (s, 3H), 3.85 (s, 3H) | δ 185.77, 170.00, 163.97, 137.29, 137.14, 56.40, 55.68 | δ 6.96 (d, *J* = 15.9 Hz, 1H), 6.32 (d, *J* = 15.7 Hz, 1H), 3.82 (s, 3H), 3.78 (s, 3H) | δ 173.95, 170.94, 147.04, 125.94, 94.63, 56.40, 55.23 | - | - |
| **1d^a^** | δ 7.75 (d, *J* = 16.0 Hz, 1H), 7.19 (d, *J* = 16.0 Hz, 1H), 6.63 (brs, 1H), 3.86 (s, 3H) | δ 183.06, 165.05, 159.42, 137.43, 132.38, 52.94 | - | - | - | - |
| **1d^e^** | δ 7.27 (d, *J* = 16.2 Hz, 1H), 6.84 (d, *J* = 16.2 Hz, 1H), 3.84 (s, 3H) | δ 197.45, 172.26, 170.28, 139.18, 137.24, 55.64 | δ 6.98 (d, *J* = 15.7 Hz, 1H), 6.26 (d, *J* = 15.7 Hz, 1H), 3.77 (s, 3H) | δ 178.10, 171.57, 150.15, 124.17, 95.82, 55.08 | - | - |
| **5b^a^** | δ 3.89 (s, 3H), 3.69 (s, 3H), 3.17 (t, *J* = 6.5 Hz, 2H), 2.69 (t, *J* = 6.5 Hz, 2H). | δ 192.39, 172.55, 161.06, 53.24, 52.18, 34.33, 27.55. | - | - | - | - |
| **5b^e^** | δ 3.89 (s, 1H), 3.69 (s, 1H), 3.24 (t, *J* = 6.5 Hz, 1H), 2.72 (t, *J* = 6.4 Hz, 1H), | δ 197.08, 178.17, 163.83, 56.16, 55.14, 36.65, 30.06. | δ 3.80 (s, 3H), 3.68 (s, 1H), 2.48 (t, *J* = 7.5 Hz, 2H), 2.18 (t, *J* = 7.5 Hz, 2H). | δ 178.69, 175.52, 96.66, 56.00, 55.05, 36.17, 30.91. | - | - |

^a^Spectra recorded in CDCl_3_ (referenced to TMS). ^b^Spectra recorded in DMSO-*d*_6_ (referenced to DMSO or TMS). ^c^Spectra recorded in 2:1 acetonitrile-*d*_3_ : D_2_O (referenced to acetonitrile). ^d^Spectra recorded in 2:1 DMSO-*d*_6_ : D_2_O (referenced to DMSO or TMS). ^e^Spectra recorded in D_2_O (referenced to DSS).

**Table S4.** ^1^H and ^13^C NMR of Criegee intermediates of **1a-c** and **5b** at 22°C.

| 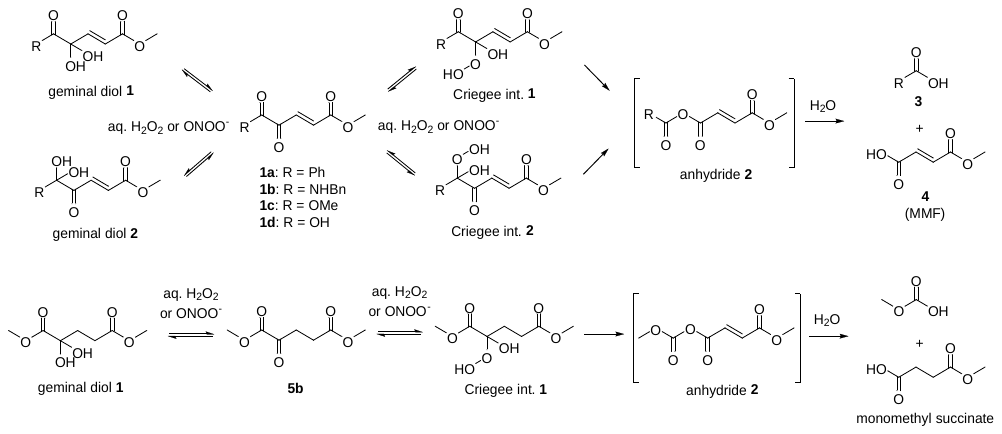 | | | | |
| --- | --- | --- | --- | --- |
| **Compound** | **Criegee Intermediate 1** | | **Criegee Intermediate 2** | |
|  | **^1^H NMR (500 MHz)** | **^13^C NMR (125 MHz)** | **^1^H NMR (500 MHz)** | **^13^C NMR (125 MHz)** |
| **1a^a^** | Olefinic peaks δ 6.72 (d, *J* = 15.8 Hz, 1H) other peak masked | - | Olefinic peaks δ 6.94 (d, *J* = 15.8 Hz, 1H), 6.24 (d, *J* = 15.7 Hz, 1H). | - |
| **1b^a^** | δ 7.44 – 7.35 (m, 5H), 6.77 (d, *J* = 15.8 Hz, 1H), 6.26 (d, *J* = 15.8 Hz, 1H), 5.28 - 5.22 (m, 2H), 3.72 (s, 3H). | δ 168.49, 167.12, 141.97, 136.21, 129.87, 129.67, 129.05, 126.28, 100.84, 68.64, 53.37. | - | - |
| **1c^a^** | δ 6.73 (d, *J* = 15.8 Hz, 1H), 6.27 (d, *J* = 15.8 Hz, 1H), 3.75 (s, 3H), 3.69 (s, 3H). | δ 168.49, 166.50, 139.98, 125.58, 99.94, 53.34, 52.08 | - | - |
| **1c^b^** | δ 6.86 (d, *J* = 15.8 Hz, 1H), 6.39 (d, *J* = 15.9 Hz, 1H), 3.86 (s, 3H), 3.78 (s, 3H). | δ 171.64, 170.32, 142.28, 128.79, 103.00, 56.61, 55.32. | - | - |
| **5b^b^** | δ 3.84 (s, 3H), 3.68 (s, 3H), 2.56-2.44 (m, 2H), 2.20 (dt, *J* = 14.7, 7.4 Hz, 1H), 2.10 (dt, *J* = 14.7, 7.4 Hz, 1H). | δ 178.22, 173.55, 104.96, 56.14, 55.09, 32.41, 30.55 | - | - |

^a^Spectra recorded in 2:1 acetonitrile-*d*_3_ : D_2_O (referenced to acetonitrile). ^b^Spectra recorded in D_2_O (referenced to DSS).

**Table S5.** ^1^H NMR monitoring of reaction rate with excess hydrogen peroxide at 22°C.

| 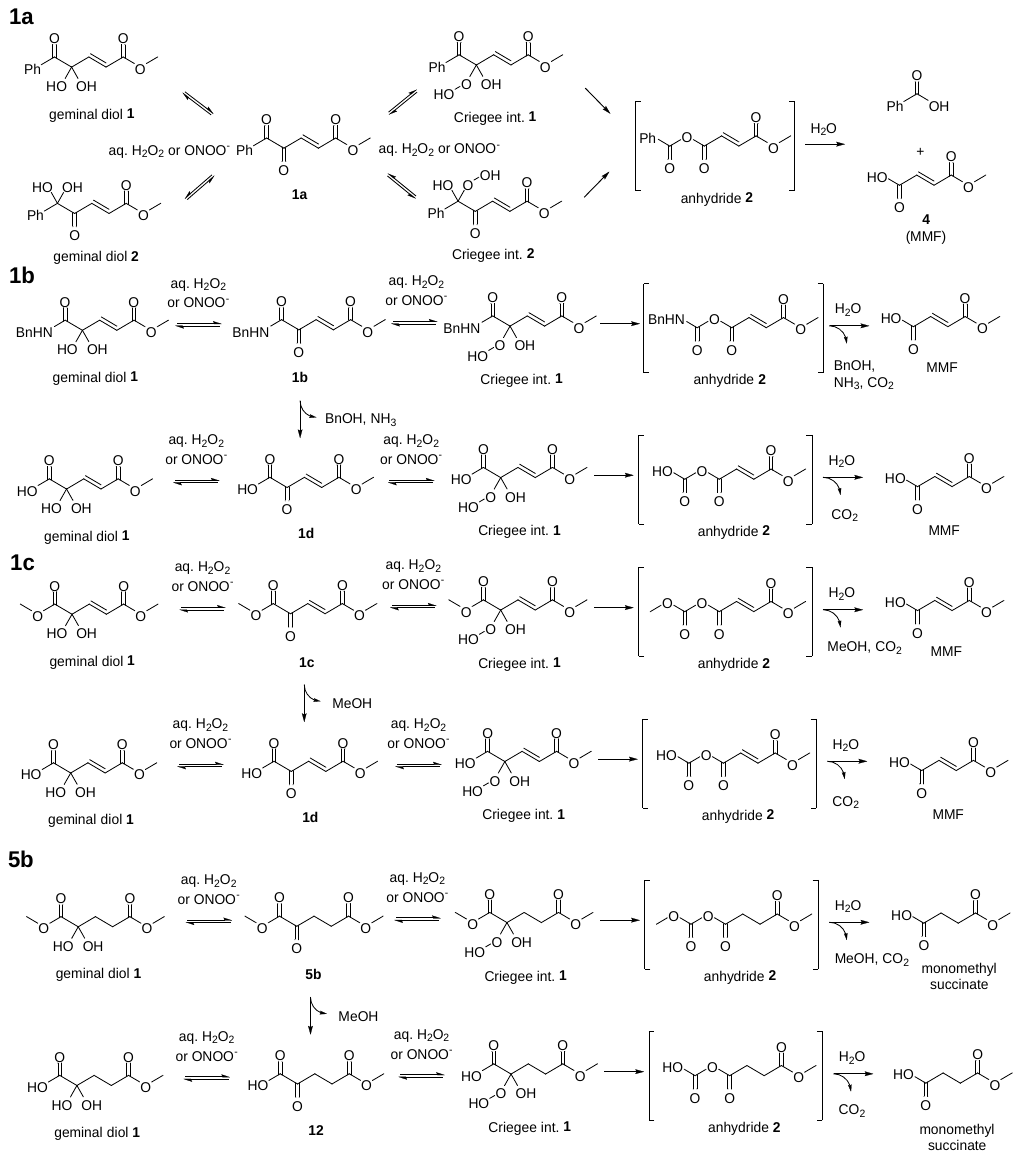 | | |
| --- | --- | --- |
| **Compound** | **Reaction half-life (t_1/2_)** | **Rate constant (k)** |
| **1a^a^** | 16 hr | 1.22 x 10^-5^ s^-1^ |
| **1b^a^** | 0.5% product formed after 7 days* | |
| **1c^a^** | 2.5% product formed after 7 days* | |
| **1c^b^** | 24.8 days* | 3.23 x 10^-7^ s^-1^ |
| **1c^c^** | 3.5 minutes | 3.30 x 10^-3^ s^-1^ |
| **1d^b^** | 4.5 minutes | 2.57 x 10^-3^ s^-1^ |
| **5b^b^** | 46.8 days* | 1.65 x 10^-7^ s^-1^ |
| **5b^c^** | 6 hours | 3.29 x 10^-5^ s^-1^ |

^a^Spectra recorded in 2:1 acetonitrile-*d*_3_:D_2_O. ^b^Spectra recorded in D_2_O. ^c^Spectra recorded in 100 mM phosphate buffer (pH7.4). *No hydrolysis to **1d** or **12** is observed in NMR reactions in D_2_O.

**Table S6.** Hydrolysis of **1c** and **5b** in D_2_O and buffer at 22°C.

| 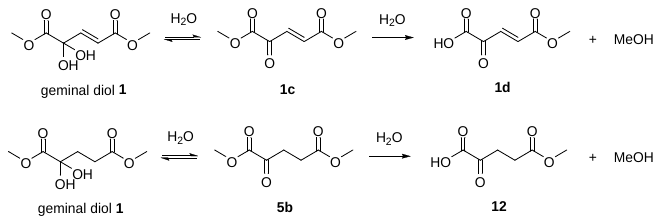 | | |
| --- | --- | --- |
| **Compound** | **Reaction half-life (t_1/2_)** | **Rate constant (k)** |
| **1c^a^** | 9.7 days | 8.25 x 10^-7^ s^-1^ |
| **1c^b^** | 20 minutes | 5.78 x 10^-4^ s^-1^ |
| **5b^a^** | 20.6 days | 3.90 x 10^-7^ s^-1^ |
| **5b^b^** | 17 minutes | 6.80 x 10^-4^ s^-1^ |

^a^Spectra recorded in D_2_O. ^b^Spectra recorded in 100 mM phosphate buffer (pH 7.4).

**Table S7. PCR primer sequences.**

| **Gene name** | **Species** | **Sequences** |
| --- | --- | --- |
| *Gapdh* | Human | F: GTCTCCTCTGACTTCAACAGCG  R: ACCACCCTGTTGCTGTAGCCAA |
| *Gclm* | Human | F: GGAACCTGCTGAACTGGGG  R: CCCTGACCAAATCTGGGTTGA |
| *Hmox1* | Human | F: CTTTCAGAAGGGCCAGGTGA  R: GTAGACAGGGGCGAAGACTG |
| *Sod1* | Human | F: ACAAAGATGGTGTGGCCGAT  R: AACGACTTCCAGCGTTTCCT |
| *Cat* | Mouse | F: CAGATGGAGAGGCAGTCTATTG  R: AAAGATCTCGGAGGCCATAATC |
| *Gapdh* | Mouse | F: TCTCCCTCACAATTTCCATCC  R: GGGTGCAGCGAACTTTATTG |
| *Gstm1* | Mouse | F: CCTCAAGAAGATCTCTGCCTAC  R: GCAAGGGCCTACTTGTTACT |
| *Hmox1* | Mouse | F: GACATGGCCTTCTGGTATGG  R: CTCGTGGAGACGCTTTACATAG |
| *Sod1* | Mouse | F: ACAATGGTGGTCCATGAGAAA  R: GTTTACTGCGCAATCCCAATC |
