## Supplementary Information for "Drug targeting to sites of oxidative stress using the Baeyer-Villiger reaction"

**Synthesis of compounds 1a-d**

**methyl (2*E*)-4-(benzylcarbamoyl)-4-oxobut-2-enoate (1b).** To a solution of methyl 4-(benzylcarbamoyl)-4-oxobutanoate (**5a**) (1.0 equiv.) dissolved in anhydrous DCM (15 ml / 1 mmol of alpha-ketoglutaric acid derivative) was added bromine (1.5 equiv.) dropwise. The mixture was stirred at 35°C under an inert atmosphere until complete. Once complete, the volatiles were removed *in vacuo* and the crude product was used without further purification. Triethylamine (2 equiv.) was added to a solution of bromide intermediate (1 equiv.) in anhydrous THF (10 ml / 1 mmol of bromide) and the mixture stirred under an inert atmosphere, protected from light at ambient temperature until complete. Once complete the volatiles were removed *in vacuo* and the crude residue purified by column chromatography to give the title compound (173 mg, 73% yield) as a yellow solid. Characterisation data matched literature reported values ^1^. ^1^H and ^13^C NMR spectra are shown in Table S3.

**1,5-dimethyl (2*E*)-4-oxopent-2-enedioate (1c).** To a solution of dimethyl 2-oxoglutarate (**5b**) (1.0 equiv.) dissolved in anhydrous DCM (15 ml / 1 mmol of alpha-ketoglutaric acid derivative) was added bromine (1.5 equiv.) dropwise. The mixture was stirred at ambient temperature under an inert atmosphere until complete. Once complete, the volatiles were removed *in vacuo* and the crude product was used without further purification. Triethylamine (2 equiv.) was added to a solution of bromide intermediate (1 equiv.) in anhydrous THF (10 ml / 1 mmol of bromide) and the mixture stirred under an inert atmosphere, protected from light at ambient temperature until complete. Once complete the volatiles were removed *in vacuo* and the crude residue purified by column chromatography to give the title compound (1.999 g, 80% yield) as a yellow solid. Characterisation data matched authentic sample purchased from commercial source. ^1^H and ^13^C NMR spectra are shown in Table S3.

**methyl (2*E*,4*E*)-5-phenylpenta-2,4-dienoate (8).**

A suspension of *trans*-cinnamaldehyde (**6**) (1 equiv.) and methyl (triphenylphosphoranylidene)acetate (**7**) (1.1 equiv.) in toluene (1 ml / 1 mmol. of aldehyde) was placed in a pressure vessel equipped with a magnetic stir bar. The vial was sealed and placed in an oil bath at 150 °C for 10 min. The reaction mixture was cooled and then transferred to a round-bottom flask and the volatiles removed *in vacuo*. Hexane or a mixture of 10% ethyl acetate in hexane was added and the mixture stirred for 10 minutes then filtered through Celite® to remove the majority of the triphenylphosphine oxide by-product. The filtrate was concentrated in vacuo the crude residue purified by column chromatography to give the title compound (1.541 g, 82% yield) as a colourless solid. Characterisation data matched literature reported values ^2^.

**methyl (2*E*)-4,5-dihydroxy-5-phenylpent-2-enoate (10).** To a 1:1 solution of t-BuOH and water (8 ml / 1 mmol. of olefin **8**) was added AD-mix β (1.4 g / 1 mmol. of olefin **8**) and the mixture was stirred at 30°C until two clear phases were visible with the lower phase bright yellow. To this mixture was added a solution of methanesulfonamide (1 equiv.) and methyl (2*E*,4*E*)-5-phenylpenta-2,4-dienoate (**8**) (1 equiv.) dissolved in a 1:1 solution of *t*-BuOH and water (2 ml / 1 mmol. of olefin) and heating at 30°C continued. Once complete sodium sulfite (1.5 g / 1 mmol of olefin **8**) was added, and the mixture stirred at ambient temperature for 30 minutes. Diethyl ether (10 ml / 1 mmol olefin **8**) was added to the reaction mixture, and after separation of the layers, the aqueous phase was further extracted with the organic diethyl ether (3 x 5 ml / 1 mmol olefin **8**). The combined organic extracts were dried (MgSO_4_), filtered and concentrated *in vacuo* to give the crude diol which was purified by column chromatography to give the title compound (403 mg, 34% yield) as a colourless oil. R*_f_* = 0.5 (50% EA in hexane); ^1^H NMR (CDCl_3_, 500 MHz) δ 7.37-7.31 (m, 5H), 6.74 (dd, *J* = 16.0, 4.0 Hz, 1H), 6.09 (dd, *J* = 16.0, 1.5 Hz, 1H), 4.55-4.53 (m, 1H), 4.41 (brs, 1H), 3.70 (s, 3H), 2.84 (d, *J* = 4.0 Hz, 1H), 2.78 (d, *J* = 2.5 Hz, 1H); ^13^C NMR (CDCl_3_, 125 MHz) δ 166.6, 145.6, 139.6, 128.7, 128.6, 126.8, 122.0, 77.0, 75.3, 51.6; IR (neat) 3425, 1705, 1660, 1495, 1449, 1437, 1391, 1311, 1277 cm^-1^.

**methyl (2E)-4,5-dioxo-5-phenylpent-2-enoate (1a).** To diol **10** (1 equiv.) that was cooled in an ice bath was added the Dess-Martin periodinane (2.1 equiv.) and the mixture stirred with a glass stirring rod until complete. Once complete the crude residue was purified by column chromatography to give the title compound (318 mg, 71%) as a yellow oil. R*_f_* = 0.55 (DCM); HRMS (ESI): calculated for C_12_H_10_O_4_ 219.0657 [M+H]^+^, found 219.0648; IR (neat) 3067, 2955, 1727, 1670, 1622, 1596, 1450, 1437, 1310 cm^-1^. ^1^H and ^13^C NMR spectra are shown in Table S3.

**(3*E*)-5-methoxy-2,5-dioxopent-3-enoic acid (1d).** **1c** (300mg, 1.74 mmol.) was dissolved in 30 ml of 100 mM phosphate buffer (pH 7.4) and stirred overnight at ambient temperature, protected from light. The solution was then diluted with 90 ml of 0.1 M HCl and then 1 M HCl was used to adjust the pH of the solution to pH 1. The mixture was then extracted with 3:1 chloroform/isopropanol 5 x 60 ml. Combined organics were dried (MgSO_4_), filtered and concentrated *in vacuo*. DCM was added to the crude residue and removed in vacuo again, this step was repeated twice more to ensure all isopropanol is removed. The crude residue was then placed under vacuum overnight, then dissolved in diethyl ether and filtered to remove the insoluble material. Volatiles were then removed in vacuo and the pure **1d** was placed under vacuum overnight, to yield **1d** (194 mg, 70% yield) as a yellow solid. IR (neat) 3536, 3344, 3084, 3067, 2967, 1770, 1732, 1687, 1452, 1440, 1370, 1314, 1263, 1217 cm^-1^; Mp 44-45°C. ^1^H and ^13^C NMR spectra are shown in Table S3.

**NMR reaction rate experiments shown in Table S5**

To **1a** (10.14 mg) in 467 μl acetonitrile-*d*_3_ and 233 μl D_2_O was added 24 μl 30% w/w hydrogen peroxide solution and the rate of MMF formation quantified by NMR spectroscopy.

| **Time (min)** | **% MMF formed** |
| --- | --- |
| 0 | 0 |
| 4 | 2 |
| 27 | 6 |
| 48 | 9 |
| 109 | 15 |
| 156 | 19 |
| 259 | 27 |
| 348 | 30 |
| 437 | 34 |
| 1143 | 56 |

MMF formation over time

| Time (min) | % MMF |
| --- | --- |
| 0 | 0 |
| 4 | 2 |
| 27 | 6 |
| 48 | 9 |
| 109 | 15 |
| 156 | 19 |
| 259 | 27 |
| 348 | 30 |
| 437 | 34 |
| 1143 | 56 |

Reaction half-life (t_1/2_): 16 hr

Rate constant (κ): 1.22 x 10^-5^ s^-1^

To **1b** (11.5 mg) in 467 μl acetonitrile-*d*_3_ and 233 μl D_2_O was added 24 μl 30% w/w hydrogen peroxide solution and the rate of MMF formation quantified by NMR spectroscopy. 0.5% product formed after 7 days.

To **1c** (8 mg) in 467 μl acetonitrile-*d*_3_ and 233 μl D_2_O was added 24 μl 30% w/w hydrogen peroxide solution and the rate of MMF formation quantified by NMR spectroscopy. 2.5% product formed after 7 days.

To **1c** (8 mg) in 700 μl D_2_O was added 24 μl 30% w/w hydrogen peroxide solution and the rate of MMF formation quantified by NMR spectroscopy.

| **Time (min)** | **% MMF formed** |
| --- | --- |
| 0 | 0 |
| 4 | 0 |
| 1222 | 1 |
| 7061 | 8.8 |
| 8721 | 10.5 |
| 19850 | 27.1 |
| 22897 | 30.4 |
| 27135 | 37.5 |
| 33260 | 46.5 |
| 37436 | 51.5 |
| 41561 | 55.2 |
| 47396 | 62 |
| 53305 | 67.9 |
| 57477 | 71.6 |

MMF formation over time

| Time (min) | % MMF |
| --- | --- |
| 0 | 0 |
| 4 | 0 |
| 1222 | 1 |
| 7061 | 8.8 |
| 8721 | 10.5 |
| 19850 | 27.1 |
| 22897 | 30.4 |
| 27135 | 37.5 |
| 33260 | 46.5 |
| 37436 | 51.5 |
| 41561 | 55.2 |
| 47396 | 62 |
| 53305 | 67.9 |
| 57477 | 71.6 |

Reaction half-life (t_1/2_): 24.8 days

Rate constant (κ): 3.23 x 10^-7^ s^-1^

The rate of MMF formation was quantified by NMR spectroscopy of a sample of **1c** (4.57 mg) in 400 μl 100 mM phosphate buffer (pH 7.4) containing 13.7 μl of 30% w/w hydrogen peroxide solution. The sample had an NMR insert to lock onto containing 200 μl of D_2_O containing DSS internal standard (200 μl of a solution of 1 mg in 700 μl of D_2_O).

| **Time (min)** | **% MMF formed** |
| --- | --- |
| 0 | 0 |
| 5 | 62 |
| 12 | 75 |

Monomethyl succinate formation over time

| Time (minutes) | % MMF |
| --- | --- |
| 0 | 0 |
| 5 | 62 |
| 12 | 75 |

Reaction half-life (t_1/2_): 3.5 minutes

Rate constant (κ): 3.30 x 10^-3^ s^-1^

To **1d** (7.35 mg) in 700 μl D_2_O was added 24 μl 30% w/w hydrogen peroxide solution and the rate of MMF formation quantified by NMR spectroscopy.

| **Time (seconds)** | **% MMF formed** |
| --- | --- |
| 0 | 0 |
| 240 | 48 |
| 840 | 90 |
| 1140 | 100 |

MMF formation over time

| Time (seconds) | % MMF |
| --- | --- |
| 0 | 0 |
| 240 | 48 |
| 840 | 90 |
| 1140 | 100 |

Reaction half-life (t_1/2_): 4.5 minutes

Rate constant (κ): 2.57 x 10^-3^ s^-1^

To **5b** (7.35 mg) in 700 μl D_2_O was added 24 μl 30% w/w hydrogen peroxide solution and the rate of monomethyl succinate (MMS) formation quantified by NMR spectroscopy.

| **Time (min)** | **% MMS formed** |
| --- | --- |
| 0 | 0 |
| 8 | 0 |
| 19 | 1 |
| 27 | 1 |
| 80 | 1 |
| 5790 | 5.5 |
| 11926 | 10 |
| 16088 | 12.4 |
| 20235 | 15 |
| 26059 | 19.4 |
| 31972 | 22.3 |
| 36144 | 25 |
| 42003 | 30 |
| 52130 | 36 |
| 70005 | 50 |

Monomethyl succinate formation over time

| Time (minutes) | % monomethyl succinate |
| --- | --- |
| 0 | 0 |
| 8 | 0 |
| 19 | 1 |
| 27 | 1 |
| 80 | 1 |
| 5790 | 5.5 |
| 11926 | 10 |
| 16088 | 12.4 |
| 20235 | 15 |
| 26059 | 19.4 |
| 31972 | 22.3 |
| 36144 | 25 |
| 42003 | 30 |
| 52130 | 36 |
| 70005 | 50 |

Reaction half-life (t_1/2_): 48.6 days

Rate constant (κ): 1.65 x 10^-7^ s^-1^

The rate of monomethyl succinate (MMS) formation was quantified by NMR spectroscopy of a sample of **5b** (4.62 mg) in 400 μl 100 mM phosphate buffer (pH 7.4) containing 13.7 μl of 30% w/w hydrogen peroxide solution. The sample had an NMR insert to lock onto containing 200 μl of D_2_O containing DSS internal standard (200 μl of a solution of 1 mg in 700 μl of D_2_O).

| **Time (min)** | **% MMS formed** |
| --- | --- |
| 0 | 0 |
| 12 | 15 |
| 33 | 25 |
| 83 | 27.5 |
| 105 | 31.5 |
| 165 | 37.9 |
| 298 | 48.2 |
| 406 | 51.7 |

Monomethyl succinate formation over time

| Time (minutes) | % MMS |
| --- | --- |
| 0 | 0 |
| 12 | 15 |
| 33 | 25 |
| 105 | 31.5 |
| 165 | 37.9 |
| 298 | 48.2 |
| 406 | 51.7 |

Reaction half-life (t_1/2_): 6 hours

Rate constant (κ): 3.29 x 10^-5^ s^-1^

**NMR reaction rate experiments shown in Table S6**

The rate of methanol formation was quantified by NMR spectroscopy of a sample of **1c** (8 mg) in 700 μl D_2_O.

| **Time (min)** | **% methanol formed** |
| --- | --- |
| 0 | 0 |
| 1245 | 3 |
| 5587 | 14.7 |
| 11,682 | 39.8 |
| 15,880 | 57.9 |
| 20,005 | 69.5 |
| 25,823 | 84 |
| 31,758 | 89.8 |
| 35,929 | 92.5 |

Methanol formation over time

| Time (minutes) | % methanol |
| --- | --- |
| 0 | 0 |
| 1245 | 3 |
| 5587 | 14.7 |
| 11682 | 39.8 |
| 15880 | 57.9 |
| 20005 | 69.5 |
| 25823 | 84 |
| 31758 | 89.8 |
| 35929 | 92.5 |

Reaction half-life (t_1/2_): 9.7 days

Rate constant (κ): 8.25 x 10^-7^ s^-1^

The rate of product (**1d** and its geminal diol) formation was quantified by NMR spectroscopy of a sample of **1c** (4.57 mg) in 400 μl 100 mM phosphate buffer (pH 7.4) with NMR insert containing 200 μl of D_2_O containing DSS internal standard (200 μl of a solution of 1 mg in 700 μl of D_2_O).

| **Time (min)** | **% product formed** |
| --- | --- |
| 0 | 0 |
| 18 | 46.7 |
| 39 | 66.2 |
| 114 | 83.9 |
| 481 | 100 |

Product formation over time

| Time (minutes) | % product |
| --- | --- |
| 0 | 0 |
| 18 | 46.7 |
| 39 | 66.2 |
| 114 | 83.9 |
| 481 | 100 |

Reaction half-life (t_1/2_): 20 minutes

Rate constant (κ): 5.78 x 10^-4^ s^-1^

The rate of methanol formation was quantified by NMR spectroscopy of a sample of **5b** (8.1 mg) in 700 μl D_2_O.

| **Time (min)** | **% methanol formed** |
| --- | --- |
| 0 | 0 |
| 4121 | 2.2 |
| 8459 | 8.9 |
| 14,561 | 19.9 |
| 18,760 | 28.8 |
| 22,885 | 36.8 |
| 28,735 | 48.5 |
| 34,637 | 58.8 |
| 38,808 | 64.8 |
| 44,632 | 73.1 |

Methanol formation over time

| Time (minutes) | % methanol |
| --- | --- |
| 0 | 0 |
| 4121 | 2.2 |
| 8459 | 8.9 |
| 14561 | 19.9 |
| 18760 | 28.8 |
| 22885 | 36.8 |
| 28735 | 48.5 |
| 34637 | 58.8 |
| 38808 | 64.8 |
| 44632 | 73.1 |

Reaction half-life (t_1/2_): 20.6 days

Rate constant (κ): 3.90 x 10^-7^ s^-1^

The rate of methanol formation was quantified by NMR spectroscopy of a sample of **5b** (4.62 mg) in 400 μl 100 mM phosphate buffer (pH 7.4) with NMR insert containing 200 μl of D_2_O containing DSS internal standard (200 μl of a solution of 1 mg in 700 μl of D_2_O).

| **Time (min)** | **% methanol formed** |
| --- | --- |
| 0 | 0 |
| 8 | 45 |
| 17 | 49.3 |
| 26 | 55.8 |
| 33 | 70.1 |
| 68 | 75.9 |
| 134 | 86.1 |
| 335 | 94.6 |
| 488 | 95.5 |

Methanol formation over time

| Time (minutes) | % methanol |
| --- | --- |
| 0 | 0 |
| 8 | 45 |
| 17 | 49.3 |
| 26 | 55.8 |
| 33 | 70.1 |
| 68 | 75.9 |
| 134 | 86.1 |
| 335 | 94.6 |
| 488 | 95.5 |

Reaction half-life (t_1/2_): 17 minutes

Rate constant (κ): 6.80 x 10^-4^ s^-1^
